# SymSim: simulating multi-faceted variability in single cell RNA sequencing

**DOI:** 10.1101/378646

**Authors:** Xiuwei Zhang, Chenling Xu, Nir Yosef

## Abstract

The abundance of new computational methods for processing and interpreting transcriptomes at a single cell level raises the need for *in-silico* platforms for evaluation and validation. Simulated datasets which resemble the properties of real datasets can aid in method development and prioritization as well as in questions in experimental design by providing an objective ground truth. Here, we present SymSim, a simulator software that explicitly models the processes that give rise to data observed in single cell RNA-Seq experiments. The components of the SymSim pipeline pertain to the three primary sources of variation in single cell RNA-Seq data: noise intrinsic to the process of transcription, extrinsic variation that is indicative of different cell states (both discrete and continuous), and technical variation due to low sensitivity and measurement noise and bias. Unlike other simulators, the parameters that govern the simulation process directly represent meaningful properties such as mRNA capture rate, the number of PCR cycles, sequencing depth, or the use of unique molecular identifiers. We demonstrate how SymSim can be used for benchmarking methods for clustering and differential expression and for examining the effects of various parameters on their performance. We also show how SymSim can be used to evaluate the number of cells required to detect a rare population and how this number deviates from the theoretical lower bound as the quality of the data decreases. SymSim is publicly available as an R package and allows users to simulate datasets with desired properties or matched with experimental data.

---

The advent of single cell RNA sequencing has led to a surge of computational and statistical methods for a range of analysis tasks. Some of the methods or the tasks that they perform have originated from bulk sequencing analysis, while others address opportunities (e.g., identification of new cell states ^1, 2^) or technical limitations (e.g., limited sensitivity ^3, 4^) that are idiosyncratic to single cell genomics ^5, 6^. While these computational methods are often based on reasonable assumptions it is difficult to compare them to each other and assess their performance without gold standards. One approach to address this is through simulations ^7–10^.

Existing simulation strategies (summarized by Zappia *et al* ^11^) rely primarily on fitting distributional models to observed data and then drawing from these distributions. While the resulting models provide a good fit to observed data, their parameters are often abstract and do not directly correspond to the actual processes that gave rise to the observations. This leaves an important unaddressed problem in designing and using a simulator: the need to modulate and then study the effects of specific aspects of the underlying physical processes, such as the efficiency of mRNA capture, the extent of amplification bias (e.g., by changing the number of PCR cycles, or by using unique molecular identifiers [UMI]), and the extent of transcriptional bursting. To address this, we present SymSim (**Sy**nthetic model of **m**ultiple variability factors for **Sim**ulation), a software for simulation of single cell RNA-Seq data. SymSim explicitly models three of the main sources of variation that govern single cell expression patterns ^2^: allele intrinsic variation, extrinsic variation, and technical factors (Figure 1). SymSim provides the users with “knobs” to control various parameters at these three levels. First, we generate “true” numbers of molecules using a kinetic model which allows us to adjust allele intrinsic variation and the extent of burst effect; second, we provide an intuitive interface to simulate a subpopulation structure, either discrete or along a continuum, through specification of cluster-trees, which define a low dimensional manifold from which the transcriptional kinetics is determined for every gene and every cell; third, we simulate the main stages of the library preparation process and let users control the amount of variation stemming from these steps, such as capture efficiency, amplification bias, varying sequencing depth and batch effect. Importantly, through this modeling scheme, SymSim recapitulates properties of the data (e.g., high abundance of zeros or increased noise in non-UMI protocols) without the need to explicitly force them as factors in a distributional model.

**Figure 1.**
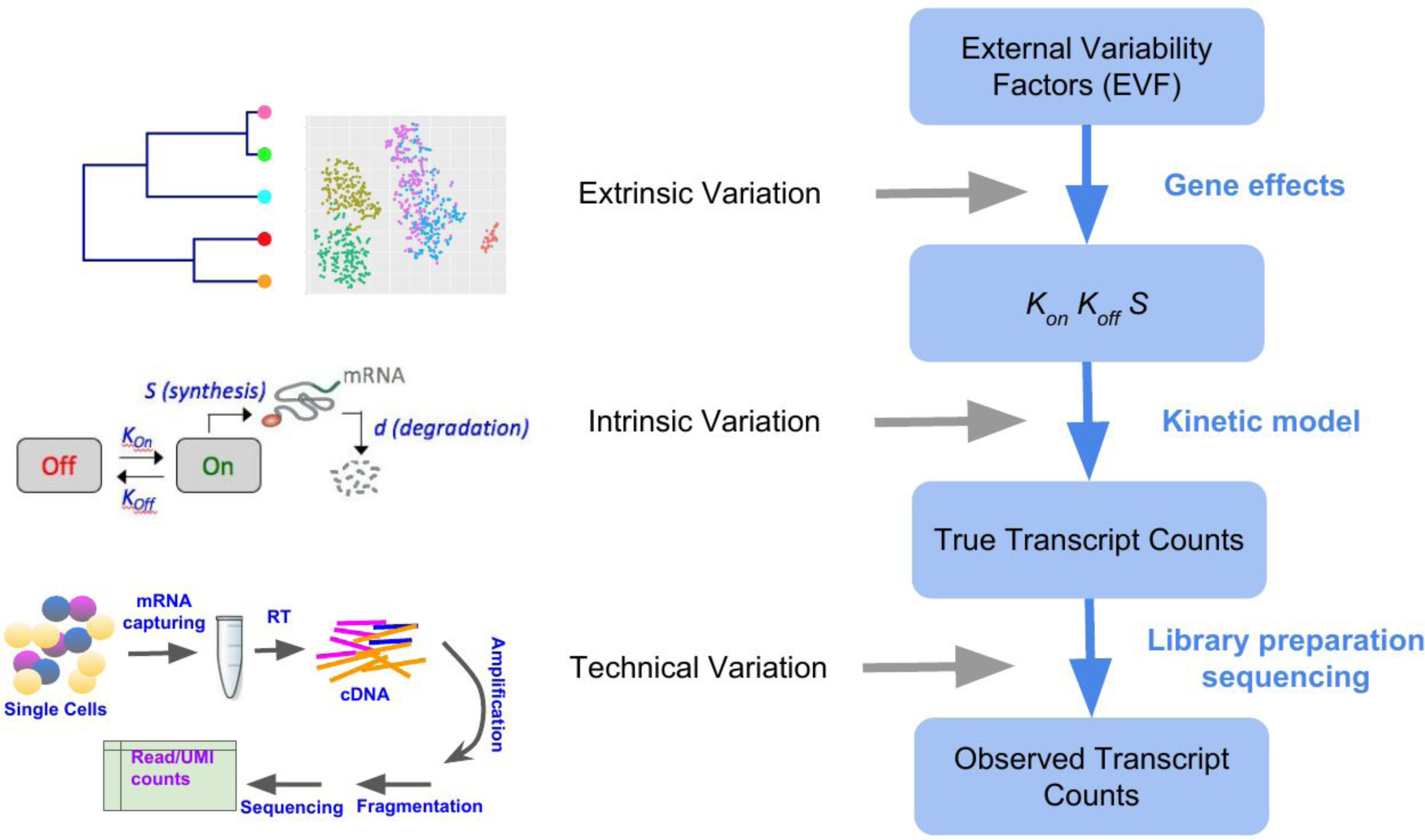
Overview of SymSim. The *true transcript counts*, which are the number of molecules for each transcript in each cell at the time of analysis, are generated through the classical promoter kinetic model with parameters: promoter *on* rate (*k_on_*), *off* rate (*k_off_*) and RNA synthesis rate (*s*). The values of the kinetic parameters are determined by the product of gene-specific coefficients (termed *gene effects*) and cell-specific coefficients. The latter set of coefficients is termed extrinsic variability factors (EVF), and it is indicative of the cell state. The expected value of each EVFs is determined in accordance to the position of the cell in a user-defined tree structure. The tree dictates the structure of the resulting cell-cell similarity map (which can be either discrete or continuous) since the distance between any two cells in the tree is proportional to the expected distance between their EVF values. For homogenous populations (represented by a single location in the tree), the EVFs are drawn *iid* from a distribution whose mean is the expected EVF value and variance is provided by the user. From the true transcript counts we explicitly simulate the key experimental steps of library preparation and sequencing, and obtain *observed counts*, which are read counts for full length mRNA sequencing protocols, and UMI counts, otherwise.

We demonstrate the utility of SymSim in two types of applications. In the first example, we use it to evaluate the performance of algorithms. We focus on the tasks of clustering and differential expression and test a number of methods under different simulation settings of biological separability and technical noise. In the second example, we use SymSim for the purpose of experimental design, focusing on the question of how many cells should one sequence to identify a certain subpopulation. The SymSim R package and an accompanying vignette are available at https://github.com/YosefLab/SymSim.

## Results

### The first knob: allele intrinsic variation

The first knob for controlling the simulation allows us to adjust the extent to which the infrequency of bursts of transcription adds variability to an otherwise homogenous population of cells. We use the widely accepted two-state kinetic model, in which the promoter switches between an *on* and an *off* states with certain probabilities ^12, 13^. We use the notation *k_on_* to represent the rate at which a gene becomes active, *k_off_* the rate of the gene becoming inactive, *s* the transcription rate, and *d* the mRNA degradation rate. For simplicity, and following previous work, we fix *d* to constant value of 1 ^12, 14^ and consider the other three parameters relative to *d* (thus becoming independent of time). Since RNA sequencing provides a single snapshot of the transcriptional process, we resort to assuming that the cells are at a steady state, and thus that the resulting single-cell measurements are drawn from the stationary distribution of the two-state kinetic model. Since *d* is fixed, we are able to express the stationary distribution for each gene analytically using a Beta-Poisson mixture (Kim and Marioni 2013) (Methods).

The values of the kinetic parameters (*k_on_*, *k_off_* and *s*) for each gene in each cell are first calculated using a product of cell-specific and gene-specific factors, then adjusted by the parameter distributions estimated from experimental data (Figure 2a, Methods). Specifically, each cell is assigned with three low-dimensional vectors (in this section, we used dimension 10; different values can be set by the user), one for each kinetic parameter. Similarly, each gene is associated with three low-dimensional vectors of the same dimension, which we term *gene effect vectors*. The value of each parameter is determined by the dot product of the two respective vectors (Figure 2a).

**Figure 2.**
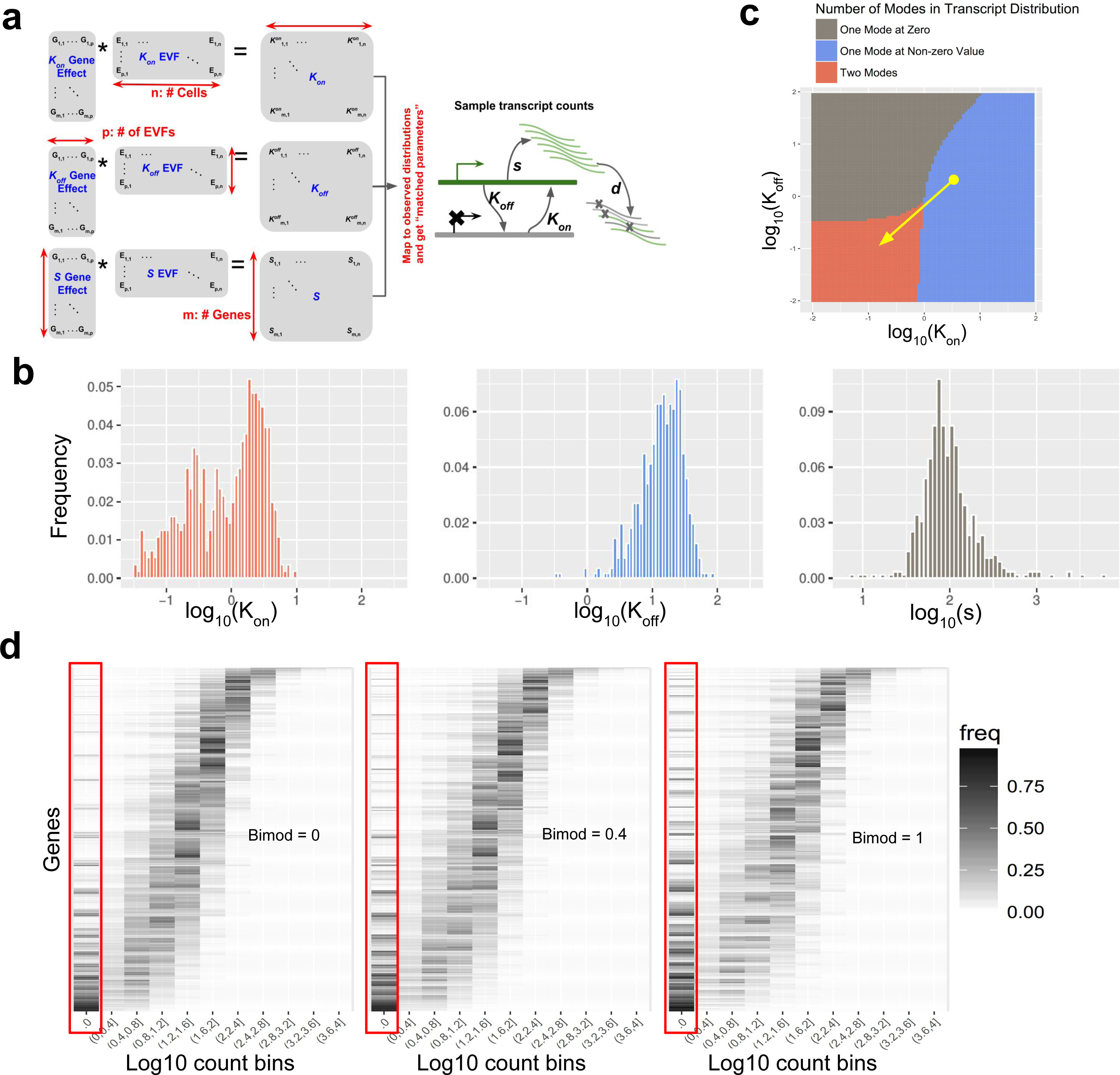
Intrinsic variation. (**a**) A diagram of how gene and cell specific kinetic parameters are simulated from cell-specific EVF and gene-specific gene effect vectors, and how the kinetic parameters are used in a model of transcription. Each cell has a separate EVF vector for *K_on_*, *K_off_* and *S*. Each parameter is generated through two steps: first, for each gene in each cell, we take the dot product of the corresponding EVF and gene effect vectors. Second, the dot product values are mapped to distributions of parameters estimated from experimental data. The “matched parameters” are used to generate true transcript counts (see Methods). (**b**) The distribution of *k_on_, k_off_* and *s* inferred from a subpopulation of oligodendrocytes after imputation by scVI that is used for the simulation. (**c**) A heatmap showing the effect of parameter *k_on_* and *k_off_* on the number of modes in transcript counts. The value of s is fixed to 10 in this plot. The red area with low *k_on_* and *k_off_* have one zero mode and one non-zero mode. The gray area with low *k_on_* and high *k_off_* has only one zero mode, and the blue area with high *k_on_* and low *k_off_* have one non-zero mode. The yellow arrow shows how the parameter *Bimod* can modify the amount of bimodality in the transcript count distribution. (**d**) Histogram heatmaps of transcript count distribution of the true simulated counts with varying values of *Bimod*, showing that increasing *Bimod* increases the zero-components of transcript counts and the number of bimodal genes. In these heatmaps, each row corresponds to a gene, each column corresponds to a level of expression, and the color intensity is proportional to the number of cells that express the respective gene at the respective expression level.

The coordinates of a cell’s vectors represent factors of cell to cell variability that are extrinsic to the noise generated intrinsically by the process of transcription (which we model by drawing from the stationary distribution above). These values, which we term *extrinsic variability factors* (EVF) represent a low dimension manifold on which the cells lie and can be interpreted as concentrations of key proteins, morphological properties, microenvironment and more. When simulating a homogeneous population, the EVFs of the cells are drawn from a normal distribution with a fixed mean of 1 and a standard deviation σ. σ is the *within-population variability* parameter and can be set by the user (for the results in this section σ is set to 0.5).

The coordinates of the gene effect vectors can be interpreted as the dependence of its kinetics on the levels of EVFs. For instance, a positive value means that higher concentration of the corresponding EVF can give rise to a higher *on* rate of a certain promoter (if the EVF and gene effect vectors are both for parameter *k_on_*). The gene effect values are first drawn independently from a standard normal distribution. We then replace each gene effect with a value of zero with probability η, thus ensuring that every gene is only affected by a small subset of EVFs. The *sparseness parameter* η can be set by the user; in this paper we set it to a fixed value of 0.7.

To ensure that the parameters used for simulation fall into realistic ranges, we estimate the distribution of kinetic parameters of genes from real data (Methods). The estimation is done by fitting a Beta-Poisson distribution to imputed experimental data. As our reference, we used a UMI based dataset of 3,005 cortex cells by Zeisel *et al* ^15^ and a non-UMI based dataset of 130 IL17-expressing T helper cells (Th17) by Gaublomme *et al* ^16^ (See Methods for further details on the experimental data). For the purpose of this analysis, the UMI based data was imputed using scVI ^4^ and the non-UMI data was imputed using ZINB-WaVE ^17^ (as scVI is only applicable to large data sets). We performed the parameter estimation separately in each of the three largest clusters in the cortex dataset (each cluster is assumed to represent a relatively homogenous subpopulation), and on the entire T cell data (a single condition, which did not contain obvious clusters) and obtained similar distribution ranges (Figures 2b and S1b). These ranges are also in line with observations from other experiments, using smFISH ^18–25^ or transcription inhibition based ^25^ methods to measure kinetic parameters (Methods and Supplementary Material Section 1). Importantly, the goal of this analysis is not to estimate the true parameter values for every gene in the reference data sets (which may not be identifiable), but rather to identify the range of plausible parameter values, to be used for simulation. To this end, SymSim applies a quantile approach to map the simulated parameter values that resulted from the dot product of the EVF and gene effect vectors to the distribution of the estimated parameters (Methods).

An intriguing question in the analysis of single cell RNA-seq data is the extent to which the conclusion drawn from the data (e.g., stratification into subpopulations) may be confounded by transcriptional bursting and transcriptional noise. SymSim provides a way to explore this. We first note that modality ^13, 26^ and extent of the intrinsic noise ^13^ in the expression of a gene in a homogenous population of cells (*i.e*., cells with similar EVFs) can vary for the different ranges of *k_on_*, *k_off_* and *s*. Specifically, one can distinguish the following three types of gene-expression distributions by the number of inflection points in the smoothed density function: unimodal with highest frequency at 0 (no inflection point), unimodal with highest frequency at non-zero value (one inflection point), and bimodal (two inflection points). Figure 2c shows the number of inflection points for different configurations of *k_on_* and *k_off_* with given *s*=10. This gives a clear correspondence between kinetic parameter configurations and types of gene-expression distributions. For example, when *s* is relatively large, we obtain bimodal distributions when *k_on_* and *k_off_* are smaller than 1.

These results thus guide us in tuning kinetic parameters to obtain desired gene-expression distributions to simulate. Specifically, we focus on adjustment of the bimodality of the distribution, which can lead to large, yet transient fluctuations in mRNA concentration at the same cell over time, thus potentially misleading methods for cell state annotation and differential expression. To increase the overall extent of bimodality in the data, we divide (decrease) all *k_on_* and *k_off_* values by 10*^Bimod^* (Figure 2c, yellow arrow). The parameter *Bimod* can take value from 0 to 1. This way, other properties such as burst frequency (*k_on_/(k_on_+k_off_)*) and synthesis rate (*s*) remain the same. In Figure 2d we show the effect of varying the *Bimod* parameter on gene-expression distribution in a simulated homogenous population. Expectedly, as *Bimod* increases, so does the number of bimodal genes, as well as the average Fano factor (Figure S1a).

### The second knob: extrinsic variation via extrinsic variability factors (EVFs)

While the first knob focuses on variation within a homogeneous set of cells, the second knob allows the user to simulate multiple, different cell states. This added complexity is achieved by setting different EVF values for different cells, in a way that allows users to control cellular heterogeneity and generate discrete sub-populations or continuous trajectories. To this end, SymSim represents the desired structure of cell states using a tree (which can be specified by the user), where every subpopulation (in the discrete mode) or every cell (in the continuous mode) is assigned with a position along the tree. Different positions in the tree correspond to different expected EVF values, and the expected absolute difference between the value of an EVF of any two cells is linearly proportional to the square root of their distance in the tree (Supplementary Material Section 2).

When SymSim is applied in a discrete mode, the cells are sampled from the leaves of the tree. The set of cells that are assigned to the same leaf in the tree form a subpopulation, and their EVF values are drawn from the same distribution. As above, we draw these EVF from a normal distribution, where the mean is determined by the position in the tree and the standard deviation is defined by the parameter σ. When SymSim is applied in a continuous mode, the cells are positioned along the edges of the tree with a small step size (which is determined by branch lengths and number of cells; Methods). The EVF values are then drawn from a normal distribution where the mean is determined by the position in the tree, and the standard deviation is defined by σ (Figure 3a).

**Figure 3.**
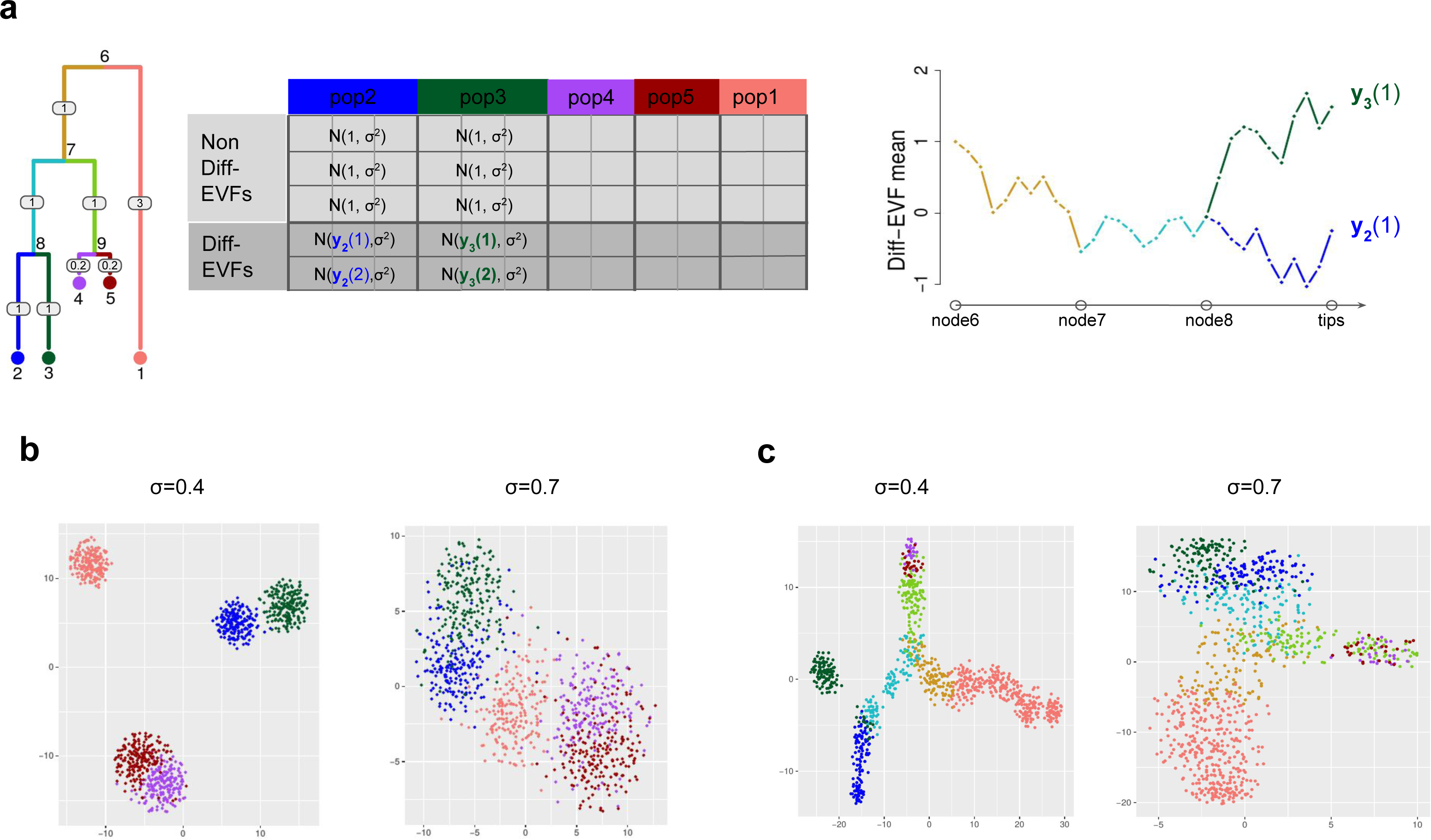
Extrinsic variation. (**a**) Illustration of generating a diverse set of cell states with SymSim. The tree represents the relationship between cells. The numbers on the edges are branch lengths; the node numbers indicate the ID of the respective subpopulation (each subpopulation is represented by a single position [leaf] in the tree). The matrix to the right depicts the derivation of EVF values. Each row corresponds to an EVF (only two are Diff-EVF), each column corresponds to a position in the tree, and the content specifies the distribution from which the EVF values are drawn. We use the notation *y_a_(b)* to represent the expected value of EVF *b* in position *a* in the tree. The rightmost plot depicts the derivation of these expected values with Brownian motion. We use subpopulations 2 and 3 as examples for both discrete cases (sampling only cells within the subpopulations) or continuous (sampling cells along the trajectories from the root ‘progenitor’ state [node 6] to the two ‘target’ subpopulations [nodes 2 and 3]). (**b**) tSNE plots of five discrete populations generated from the tree structure shown in (a). Different values of σ give rise to different heterogeneity of each population. (**c**) tSNE plots of continuous populations generated from the same tree. The colors corresponds to the colors on branches in the tree shown in (a). When increasing σ, cells are more scattered around the main paths which follow the tree structure.

To facilitate the correspondence between EVF values and distances in the tree we use a Brownian motion procedure ^27^ (Methods; Figure 3a). Specifically, for each EVF we set the mean value at the root of the tree to a fixed number (default set root node to 1) and then perform Brownian motion along the branch. Figure 3a illustrates this process using populations 2 and 3 in the tree as an example. Notably, in the continuous mode, this formulation can give rise to a rich set of patterns of changes in gene expression from root (‘progenitor cells’) to leaves (‘target cells’), including the commonly observed impulse profile ^28, 29^ (Figure S1c-d). As an alternative, we also implemented a mode for simulating continuous data by which gene expression from root to leaves is determined explicitly by an impulse function. This might be preferable if the user would like to generate smoother changes in gene expression, or specific temporal patterns. In the following analyzes we use the Brownian motion model.

Notably, SymSim only generates a subset of EVFs from the tree, while the remaining ones are drawn from the same distribution for all subpopulations (Figure 3a). The tree-sampled subset, which we term *Diff-EVFs* (Differential EVFs) represents the conditions or factors which are different between sub-populations, and they usually account for a small proportion of all the EVFs. The number of Diff-EVFs can be set by the user. The results in this section were produced with 60 EVFs, 20% of them are Diff-EVFs.

With this formulation, users can control the extent of between-population variation by setting the branch lengths of the input tree, and combine it with a desired level of within-population variation by setting the parameter σ. Notably, both σ and the square root of branch lengths in the tree are in units of EVF values. It is therefore the case that for any two positions in the tree, the ratio of square root tree distance to σ determines the separability between the respective distributions of the values assigned to any given Diff-EVF (Supplementary Material Section 2). As illustration, Figure 3 depicts the tSNE plots of cells from the same input tree with different σ in either a discrete (Figure 3b) or continuous (Figure 3c) mode. Notably, both panels show that the tSNE plots reflect the structure of the input tree well.

### The third knob: technical variation

A large part of the variation observed in scRNA-seq data sets stems from technical sources ^30–32^. The technical confounders reflect noise, reduced sensitivity and bias that are introduced during sample processing steps such as mRNA capture, reverse transcription, PCR amplification, RNA fragmentation, and sequencing. In order to introduce realistic technical variation into our model, we explicitly simulate the major steps in the experimental procedures. We implemented two library preparation protocols: (1) full length mRNAs profiling without the use of UMIs (e.g., with a standard SmartSeq2 ^33^); and (2) Profiling only the end of the mRNA molecule with addition of UMIs (e.g., 10x Chromium ^34^). The former protocol is usually applied for a small number of cells and with a large number of reads per cell, providing full information on transcript structure ^35^. The latter is normally applied for many cells with shallower sequencing, and it is affected less by amplification and gene length biases ^30^.

The workflow of these steps are shown in Figure 4a (Methods). Starting from the simulated true mRNA content of a given cell (namely, number of transcripts per gene, sampled from the stationary distribution of the promoter kinetic model), the first step is mRNA capture, where every molecule is retained with probability 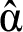. The value of the capture efficiency 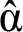 associated with each cell is drawn from a normal distribution with a mean α and standard deviation β, which can be set by the user. The second step is amplification, where in every cycle SymSim selects each available molecule with a certain probability and duplicates it. The expected amplification efficiency and the number of PCR cycles can be set by the user. As an optional step, SymSim provides the option of linear amplification (e.g., as in CEL-Seq ^36^). We do not apply this option in this manuscript. In the third step each amplified molecule is broken down into fragments, in preparation for further amplification, size selection and sequencing. The lengths of the simulated transcripts are obtained from the human reference genome, and the fragmentation is calibrated so that the average fragment length is 400bp, which is typical for RNA sequencing (Supplementary Material Section 3). Resulting fragments that are within an acceptable size range (100 to 1000 bp) are then carried on to the fourth and last step of sequencing.

**Figure 4.**
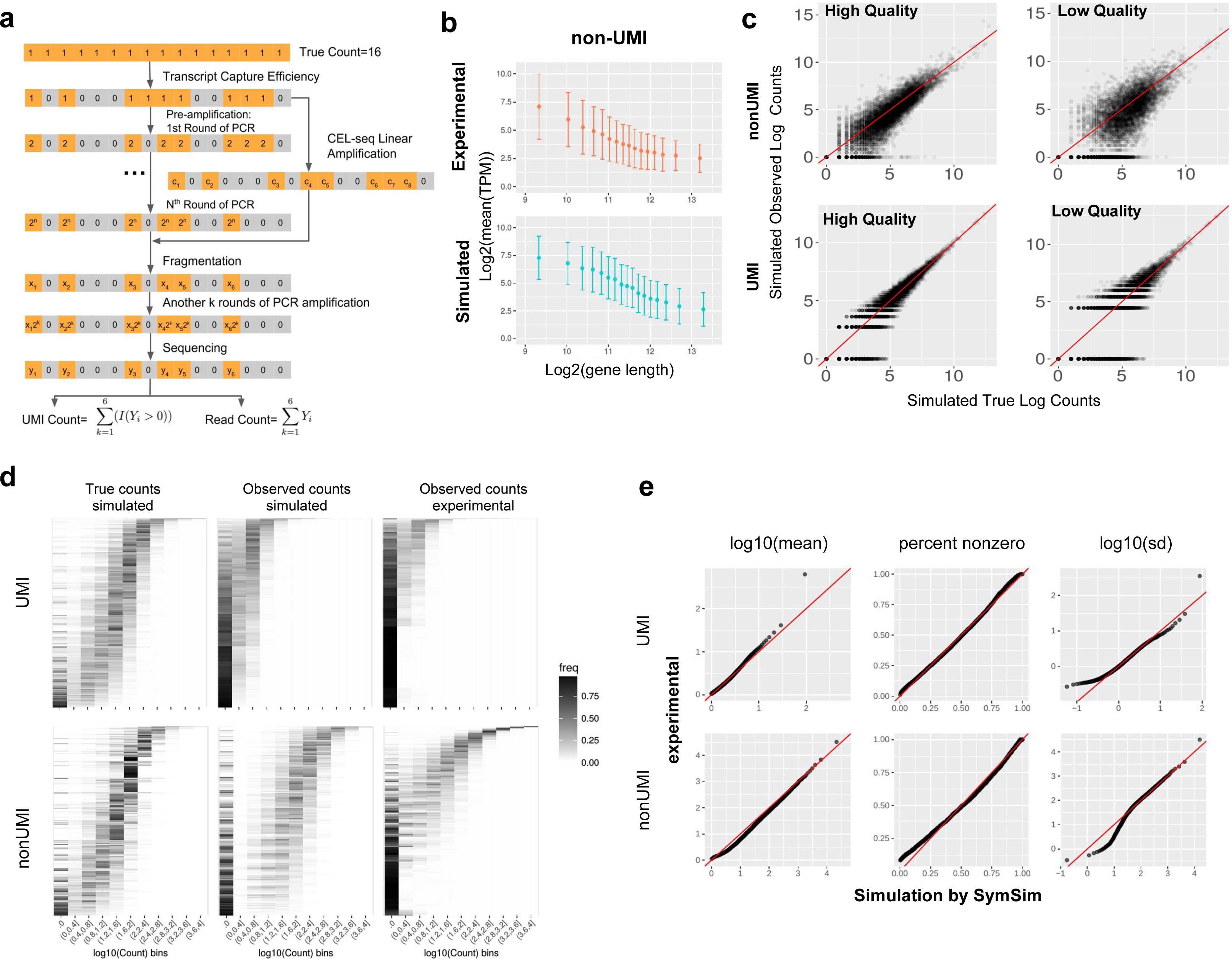
Technical variation. (**a**) A diagram showing the workflow of adding technical variation to true simulated counts. Each gray or orange square represents a molecule of the same transcript in one cell. We implement the following steps: mRNA capturing, pre-amplification (PCR or linear amplification of the cDNAs), fragmentation, amplification after fragmentation, sequencing, and calculation of UMI counts or read counts. Details of these steps can be found in Methods. (**b**) Gene length bias in both simulated and experimental data for the non-UMI protocol. (**c**) Scatter plots comparing true counts and observed counts obtained through: 1. non-UMI, “good” parameters (*α*=0.2, *MaxAmpBias*=0.1, *Depth*=1e6); 2. UMI, “good” parameters (*α*=0.2, *MaxAmpBias*=0.1, *Depth*=5e5); 3. non-UMI, “bad” parameters (*α*=0.05, *MaxAmpBias*=0.2, *Depth*=1e6); 4. UMI, “bad” parameters (*α*=0.04, *MaxAmpBias*=0.2, *Depth*=5e5). (**d**) 2D transcript counts histogram heatmap of UMI and non-UMI simulated true counts and simulated observed counts, generated with parameters which best match the input experimental counts, and histogram heatmaps of the respective experimental counts. (**e**) Q-Q plots comparing the mean, percent non-zero and standard deviation in experimental counts and simulated observed counts in non-UMI and UMI simulation. A good match is indicated by most of the dots falling close to the red line.

The number of reads per cell (namely, the number of sequenced fragments) is drawn from a normal distribution whose mean is determined by the parameter *Depth*, which, along with the respective standard deviation (*Depth_sd*) can be provided by the user. To derive the final “observed” expression values we do not account for sequencing errors, and assume that every sequenced fragment is assigned to the correct gene it originates from. For the non-UMI option, we define the raw measurement of expression as the number of reads per gene. If UMIs are used, SymSim counts every original mRNA molecule only once by collapsing all reads that originated from the same molecule. Notably, for certain depth values, the resulting distribution of number of reads per UMI is similar to the one observed in a dataset of murine cortex cells ^15^ (Figure S2a-b).

It has been previously shown that estimation of gene-expression levels from full length mRNA sequencing protocols has amplification biases related to sequence-specific properties like gene length and GC-content ^30, 37^, whereas the use of UMIs can correct these biases ^37, 38^. In particular, we have observed a negative correlation between gene length and length-normalized gene-expression in our reference non-UMI dataset (murine Th17 cells from Gaublomme et al ^16^; Figure 4b), and the same trend is reported by Phipson *et al* ^37^. To account for that, we parametrize the efficiency of the PCR amplification step using a linear model that represents gene length bias (Methods). As a result, our simulated data with a non-UMI protocol shows a similar dependence of gene-expression on gene length as in experimental data (Figure 4b, real data is from ^16^). In cases where UMIs are used, gene length effects are also modeled during amplification, but these effects are mitigated since each molecule is counted at most once. We therefore do not observe gene length bias in the UMI-based simulated data, similarly to the experimental data (Figure S2c, real data is from ^15^). Finally, we model batch effects with multiplicative factors that are gene- and batch-specific. In Figure S2d, we show the same population of cells are separated by batches. To simplify the discussion at the remainder of this paper, we assume that the data comes from a single batch.

In Figure 4c, we show the comparison between the simulated true mRNA content of one cell and the observed counts obtained with or without UMI. We consider two scenarios: the first scenario represents a study with a low technical confounding and the second one represents a highly confounded dataset. Parameters which differ between these “good” and “bad” cases in this example include capture efficiency (*α*), extent of amplification bias (*MaxAmpBias*) and sequencing depth (*Depth*). Using “bad” technical parameters introduce more noise to true counts, and compared to the non-UMI simulation the UMIs reduce technical noise. The histograms of true counts and four versions of simulated counts are shown in Figure S2e. Using quantile-quantile plots (Q-Q plots; Figure S2f) further demonstrates that UMIs help in maintaining a better representation of the true counts in the observed data.

### Fitting parameters to real data

For a given real data set, SymSim can produce observed (read or UMI) counts which have similar statistical properties to the real data (Figures 4d-e), by searching in a database of simulations obtained from a range of parameter configurations (Methods). This procedure focuses on within-population variability (Similarly to Splatter ^11^) and sets the values of eight parameters from both the first and third knobs. We test this function with the non-UMI Th17 dataset ^16^ (using all cells) and the cortex dataset ^15^ (using a subpopulation of 948 CA1 pyramidal neuron cells). See Supplementary Material Section 4 for the values of the fitted parameters.

Side by side inspection of the histograms of true mRNA levels (simulated) and observed counts (simulated and experimental), indicates that SymSim can transform the simulated ground truth (Figure 4d, left) into simulated observations (Figure 4d, middle) that match the real data for both UMI and non-UMI protocols (Figure 4d, right). For a more quantitative analysis, we generated Q-Q plots of the distributions of mean, percent-non-zero and standard deviation (SD) of genes between simulated and experimental data. Notably, we observe a certain level of inaccuracy in matching the SD at the lower ends, which can be due to lowly expressed genes. Indeed, when we filter out lowly expressed genes, the matching of SD improve substantially (Q-Q plots shown in Figure S3a). Furthemore, we conducted a similar analysis by training Splatter ^11^ with the same experimental datasets as input, and found that SymSim matches this data significantly better (Figure S3b). Finally, we inspected the relationship between mean (across all cells) and detection rate (fraction of cells in which the gene is detected) from the SymSim simulations, and observed a similar trend as in the experimental data (Figure S3c).

### Using SymSim to evaluate methods for clustering and differential expression

SymSim can be used to benchmark methods for single cell RNA-Seq data analysis as it provides both observed counts and a reference ground truth. In this section we demonstrate the utility of SymSim as tool for benchmarking methods for clustering and differential expression in a sample consisting of multiple subpopulations, using the structure depicted in Figure 3a. The design of SymSim allows us to evaluate the effect of various biological and technical confounders on the accuracy of downstream analysis. Here, we investigate the effect of total number of cells (*N*), within population variability (σ), mRNA capture rate (α) and sequencing depth (*Depth*). We also test the effect of the proportion of cells associated with the smallest sub-population of cells (*Prop*), using population 2 in the tree as our designated “rare” sub-population.

We begin by inspecting the impact of each parameter on the performance of clustering methods. To this end, we simulated observed counts using the UMI option, and traversed a grid of values for the five parameters with 18 simulation runs per configuration. The values of the remaining parameters are largely determined according to the cortex dataset ^15^ and specified in Supplementary Material Section 5.2. We tested three clustering methods: k-means based on euclidean distance of the first 10 principle components, k-means based on Euclidean distance in a nonlinear latent space learned by scVI ^4^ and SIMLR ^39^. In all cases we set the expected number of clusters to the ground truth value (n=5). The accuracy of the methods is evaluated using the adjusted Rand index (ARI; higher values indicate better performance). To inspect the effects of the various parameters on clustering performance, we performed multiple linear regression between the parameters and the ARI. The regression coefficients are shown in Figure 5a. Overall, σ appears to be the most dominant factor, and the proportion of the rare population (*Prop*) is clearly positively associated with better performance. Among the technical parameters, while α plays a role on the performance especially for the rare population, the impact of *Depth* is minor. Focusing on the dominant factors (except N, which we discuss in the next section), provides the expected results, with better accuracy as the quality of the data or the differences between subpopulations increase (Figure 5b-c). Interestingly, comparing σ=0.6, σ=0.8 and σ=1, we can tell that when σ is high enough to make the clustering challenging, further increasing σ does not yield obvious changes (Figure 5b). We observe a similar trend of saturation, inspecting increasing levels of capture efficiency (α), especially with scVI. Comparing the methods to each other, we see that scVI has the highest ARI in most cases and that PCA and SIMLR are comparable with SIMLR being slightly better when the rare population accounts for 5% of all cells and the other way around when the size of rare population increases to 10% of all cells.

**Figure 5.**
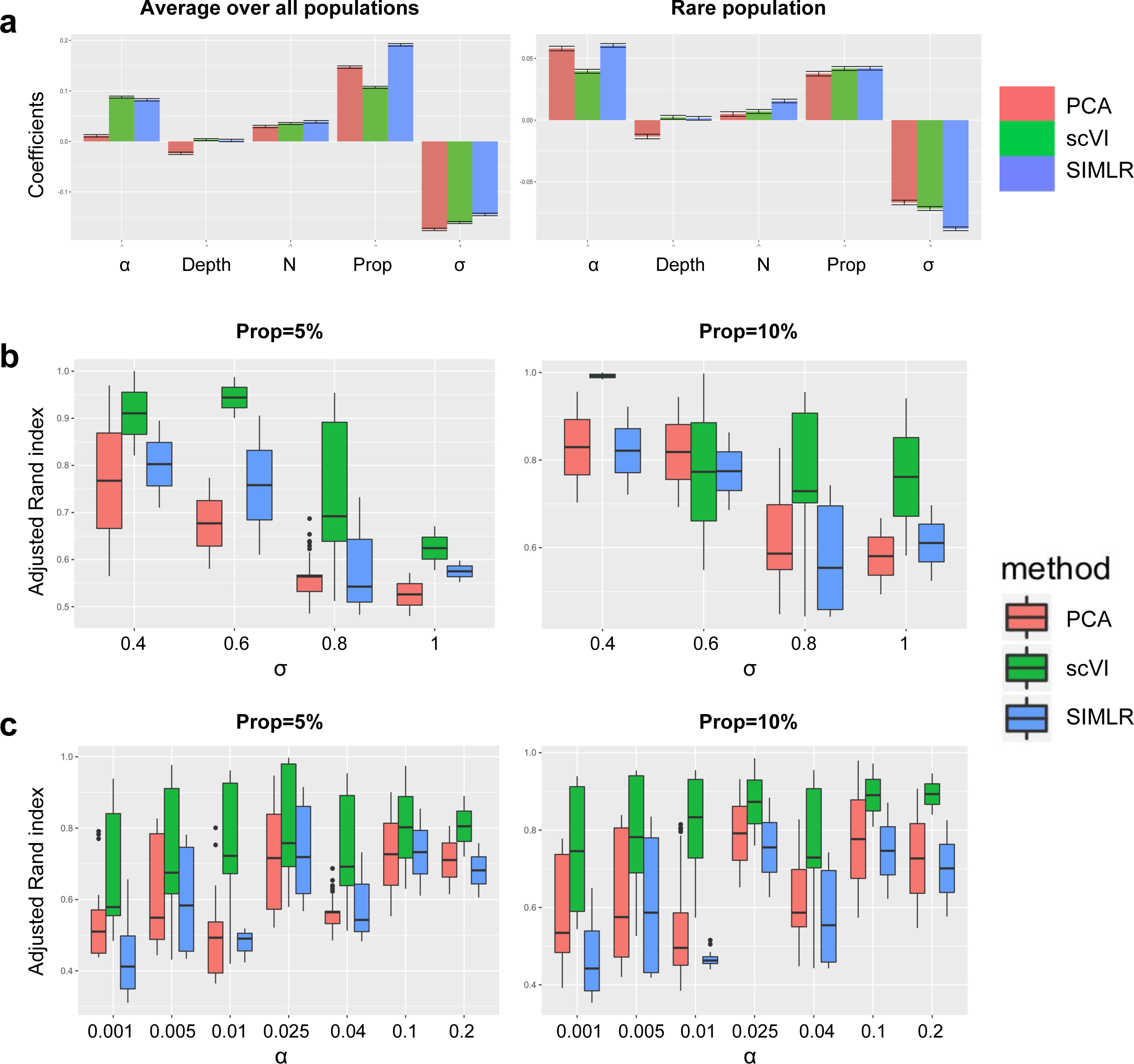
Benchmarking of clustering methods. (**a**) Coefficients of various parameters from multiple linear regression between parameters and the adjusted Rand index (ARI). In the left plot the ARI are averaged over all populations, and in the right plot the ARI is only for the rare population (population 2, with 5% of all cells). (**b**) Average ARI over all populations using the three clustering methods when changing σ (α=0.04). Left plot: the rare population accounts for 5% of all the cells; right plot: the rare population accounts for 10% of all the cells. (**c**) Average ARI over all populations using the three clustering methods when changing α (σ=0.8). Left plot: the rare population accounts for 5% of all the cells; right plot: the rare population accounts for 10% of all the cells.

Our mechanism for simulating multiple populations automatically generates differentially expressed (DE) genes between populations (in the discrete setting; Figure 3b) or along pseudotime (in the continuous setting; Figure 3c). In the following, we use SymSim to benchmark methods for detecting DE genes, focusing on the discrete setting. We use two criteria to define the ground truth set of DE genes. The first criterion is that the number of Diff-EVFs that are associated with a non-zero gene effect value (which we denote as *nDiff-EVFgene;* Figure 6a) should be larger than zero. This criterion is motivated by our model of transcription regulation: the kinetic parameters of a gene are affected by extrinsic factors, and changes to extrinsic factors might therefore lead to changes in the number of transcripts. Indeed, when we compare the true simulated gene expression values between subpopulations (i.e., before introducing technical confounders with the third knob), we get a uniform (random) distribution of p-values for genes with no Diff-EVFs, and an increasing skew as *nDiff-EVFgene* increases (Figure 6b, using Wilcoxon test); Figure S4a shows that the log fold change of gene-expression between subpopulations increases with *nDiff-EVFgene*. Nevertheless, in some cases, the actual expression values of genes with *nDiff-EVFgene>0* might not differ since the effects of different Diff-EVFs or the effects of modifying different kinetic parameters may cancel out. Differential expression might also be blurred by a high within-population variability. We therefore added an additional constraint and require that all DE genes must have a sufficiently large log fold difference in their simulated true simulated expression levels (Methods).

**Figure 6.**
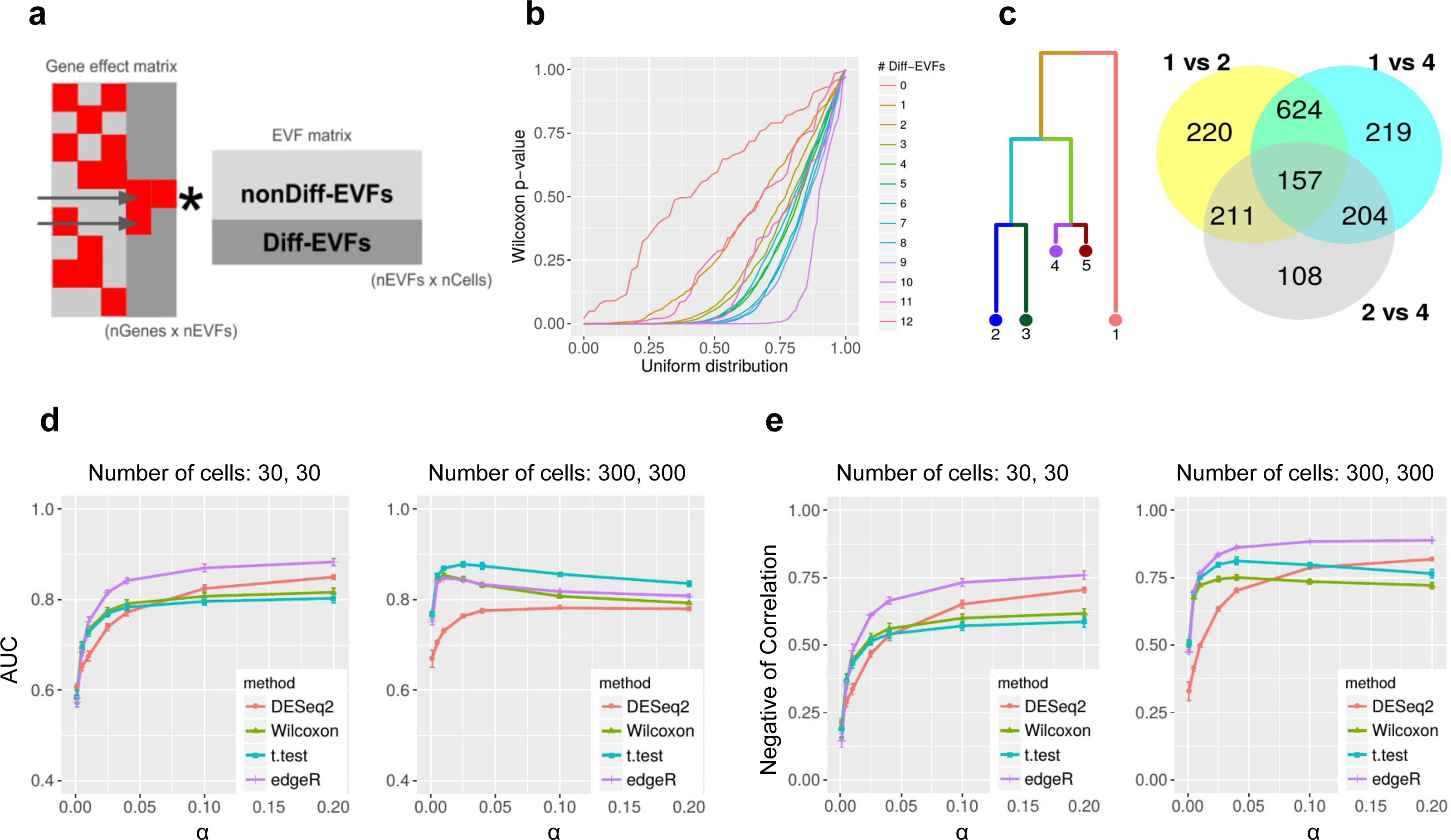
Benchmarking of DE detection methods. (**a**) Illustration of how DE genes are generated through the Diff-EVFs. Red squares in the gene effect matrix correspond to non-zero values. The two genes indicated by the arrows are DE genes by number of Diff-EVFs they have (respectively 2 and 1). (**b**) Q-Q plot comparing the p-value obtained from differential expression analysis between subpopulations 2 and 4 (using Wilcoxon test on the true simulated counts) to a uniform distribution. Genes are grouped by the number of Diff-EVFs they use and different groups are plotted in different colors. The numbers of Diff-EVFs used by genes can be thought of as the degree of DE-ness. Genes with more Diff-EVFs have p-values further diverged from uniform distribution. (**c**) Venn diagram showing that closely related populations have less DE genes between them compared to distantly related populations. We use populations 1, 2, 4 as examples: there are much more DE genes from comparison of “1 vs 2” and “1 vs 4” than “ 2 vs 4”, and DE genes from “1 vs 2” and “1 vs 4” have a big overlap. The DE genes are determined by log fold change (LFC) of true counts with criterion |LFC| > 0.8. (**d**) The AUROC (area under receiver operating characteristic curve) of detecting DE genes using four different methods from observed counts with changing capture efficiency *α* (σ=0.6). The populations under comparison are 2 and 4. Three sets of criteria were used to define the true DE genes and the final performance was the average performance from the three sets: (1) *nDiff-EVFgene*>0 and |LFC|>0.6; (2) *nDiff-EVFgene*>0 and |LFC|>0.8; (3) *nDiff-EVFgene*>0 and |LFC|>1. LFC was calculated with theoretical means from the kinetic parameters. (**e**) The negative of correlation between log fold change on theoretical mean of gene-expression and p-values obtained by a DE detection method, which changing capture efficiency α (σ=0.6). The populations under comparison are 2 and 4.

An important distinguishing feature of SymSim is that it provides an intuitive way for generating case studies for DE analysis that consist of multiple subpopulations with a predefined structure of similarity. To illustrate this, consider populations 1, 2, and 4 (Figure 6c), which form a hierarchy (2 and 4 are closer to each other and similarly distant from 1). This user-defined structure is reflected in the sizes of the sets of DE genes, obtained respectively from populations 1 vs 2 (1212 genes), 1 vs 4 (1204 genes) and 2 vs 4 (680 genes). Consistent with the hierarchy, the first two gene sets are overlapping and larger than the third one.

As an example for a benchmark study, we used four methods to detect DE genes: edgeR ^40^, DESeq2 ^41^, Wilcoxon rank-sum test and t-test on observed counts generated by various parameter settings (Methods, Supplementary Material Section 5.3). We tested the effect of the total number of cells (*N*) and mRNA capture rate (α) with 10 simulation runs per parameter configuration. We use two accuracy measures: a) AUROC (area under receiver operating characteristic curve), obtained by treating the p-values output from each method as a predictor (Figure 6d); b) negative of Spearman correlation between the p-values of each detection method and the log fold difference of the true expression levels (Figure 6e).

From Figures 6d-e, one can observe that when the numbers of cells are small (30 in each population), edgeR has the best performance while the other three methods are comparable to each other. When the numbers of cells increase to 300, the two naive methods Wilcoxon test and t-test improve in their relative performance, compared to edgeR and DESeq2. When increasing capture efficiency, all methods gain performance except for the case of AUROC with 300 cells. In that case, the drop in AUROC for some methods is caused by inflation in p-values as α increases, which results in lower specificity (Figure S4b). Notably, we noticed that the adjusted p-values from DESeq2 can have many missing entries (NAs), especially when *α* is low (and thus counts are low), and therefore we used its unadjusted p-values in Figure 6d-e. However, this assignment of NAs in practice filters out genes which do not pass a certain threshold of absolute magnitude (explained in DESeq2 vignette ^42^). To make use of this filtering, we conducted an additional analysis where we used the adjusted p-values for DESeq2 and compare it to all other methods using only the non-filtered (non NA) genes (Figure S4c). As expected, the performance of all methods (and specifically DESeq2) improves when considering only this set of genes, and converges to high values already at lower capture efficiency rates.

To summarize, we find that edgeR has the best overall performance, with the t-test rank second followed by Wilcoxon test. We note that the aim of this section is to demonstrate the use of SymSim for methods benchmarking instead of performing a comprehensive comparisons of methods. Nevertheless, our ranking is consistent with results from a recent paper which evaluated 36 methods for DE analysis with single cell RNA-Seq data ^43^.

### Experimental Design

Deciding how many cells to sequence is a decision many researchers face when designing an experiment, and the optimal number of cells to sequence highly depends on the nature of the biological system under investigation and the respective technical hurdles. A previous approach to this problem ^44^ assumes that the goal of the experiment is to identify subpopulations of cells and provides a theoretical lower bound for the problem. This bound considers the aspect of counting cells (namely, sequencing enough representative cells from each subpopulation), but it does not account for the identifiability of each subpopulation, which may be hampered by both technical and biological factors as well as the performance of clustering algorithms.

In the following we demonstrate how SymSim can be used to shed more light on this important problem. Importantly, in its current form SymSim does not use real data to model between-population variability. We therefore interpret the results in a relative manner --- how do different variability factors shift the required number of cells, compared to each other and to the theoretical lower bound. Our example focuses on a case of one rare subset, represented by cells from population 2 (using the same tree in Figure 6c; note that one can easily generalize this procedure to multiple rare subpopulations). We simulate observed counts with numbers of cells (*N*) ranging from 600 to 7000. These simulations were based on the parameters fit to the cortex dataset ^15^ with varying levels of σ and α (100 simulations per parameter configuration).

We applied the same three clustering methods as described in the previous section (*k-*means with scVI or PCA and SIMLR). We say that a given algorithm was successful in “detecting the rare population” if at least 50 cells from this set are assigned to the same cluster, and form at least 70% of the cells in that cluster. We use these labels to compute an empirical success probability *P* for each algorithm and each parameter configuration. To get an upper bound on performance that better reflects the data (rather than the algorithm), we take the maximum *P* at each configuration, and apply cubic spline smoothing (gray curves, Figure 7a-d). In each plot we also include the theoretical limit which only requires the presence of at least 50 cells from the rare subpopulation (Methods). The theoretical curve (which is independent of all parameters except N) reaches almost 1 at N=1400. Conversely, the empirical curves vary dramatically, based on parameter values. For an easy case of low within-population variability (σ=0.6) and high capture efficiency (α=0.1) the empirical upper bound curve is close to the theoretical one (Figure 7a). This curve clearly decreases when increasing the effect of either nuisance factor (Figure 7b,c). The reduction is substantially more dramatic when both nuisance factors increase (Figure 7d).

**Figure 7.**
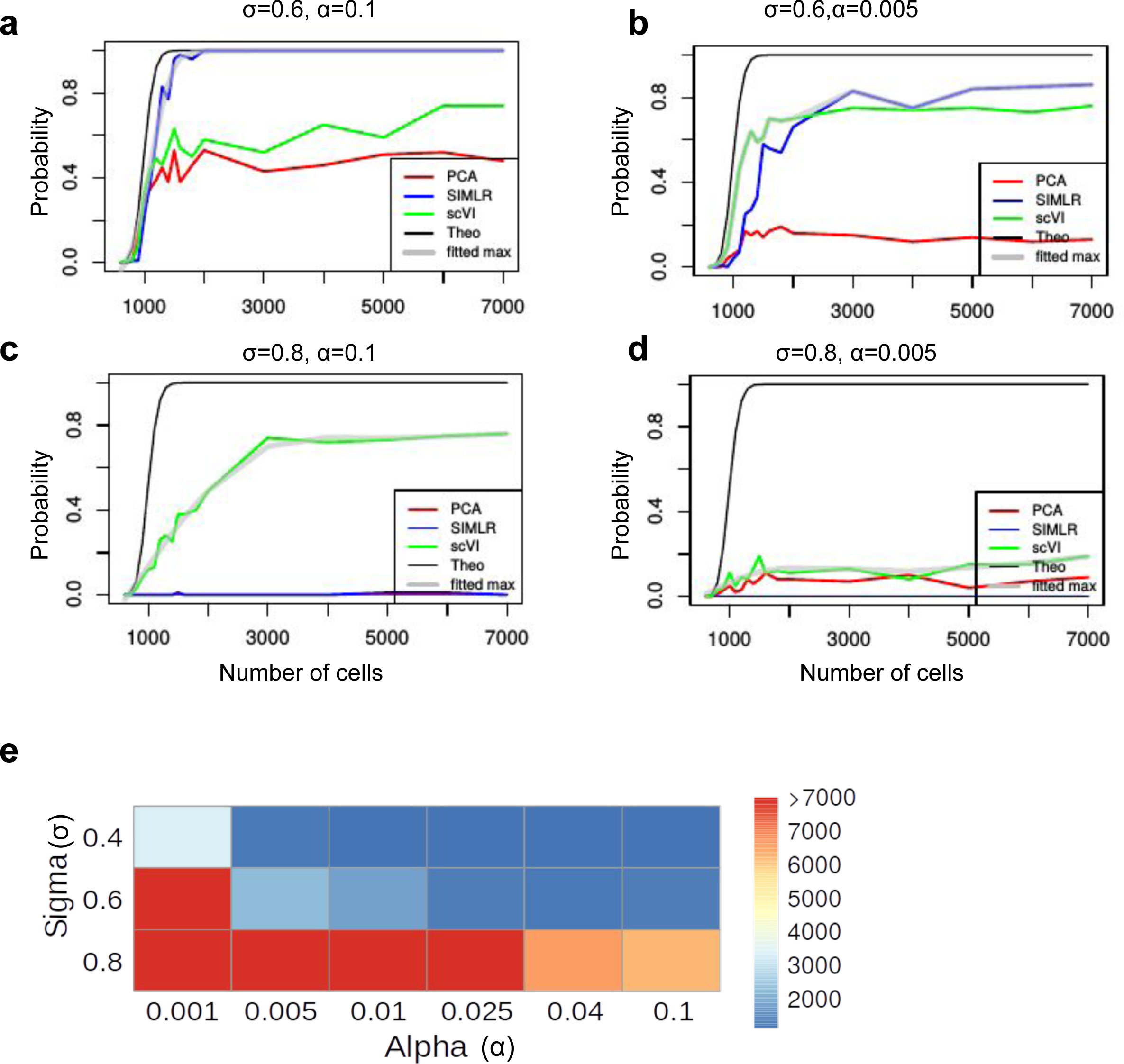
The number of cells needed to detect a rare population. We generate five populations according to the tree structure shown in Figure 3 and set population 2 as the rare population which accounts for 5% of the cells. Other populations share 95% of the cells evenly. The criteria of “detecting the rare population” is that at least 50 cells from this population are correctly detected and the precision (positive predicted value) is at least 70%. (**a-d**) The probability of detecting the rare population with when sequencing *N* (x axis) cells under different *σ* and *α* configurations, with different clustering methods. The black curve represents the theoretical probability from the binomial model, assuming that all cells sequenced are assigned correctly to the original population. The gray curve with transparency takes the maximum value at each data point from all three clustering methods with smoothing. (**e**) The heatmap shows the number of cells needed to sequence under different configurations of *σ* and *α* to detect the rare population with probability 0.75, always using the best clustering method,.

To understand the implications on the number of cells required in a given setting, we calculated how many cells are required, in each configuration, to achieve a success rate of 0.75 (*P*=0.75, Figure 7e). As expected, the resulting numbers can be much higher than the theoretical lower bound. For example, even when we have good capture efficiency (α=0.1), when the within-population variability increases (σ=0.8), we need 6225 cells, while with the theoretical curve, we need only less than 1100 cells. Considering only the binomial sampling of cells may therefore underestimate the number of cells needed for a realistic scenario, and considerations of biological and technical variations with simulators like SymSim is merited.

## Discussion

SymSim has the following features which are advantageous over existing simulators: (i) We simulate true transcript counts from a kinetic model that can be interpreted in terms of transcript synthesis rate, promoter activation and deactivation. (ii) When generating multiple discrete or continuous populations, instead of generating biological differences through directly altering the true transcript count distribution, we set Diff-EVFs, which can be interpreted as biological conditions that cause the differences between subpopulations of cells. This is a more natural and realistic way to simulate biological transcriptional differences. (iii) The EVF formulation provides an intuitive way to specify and simulate complex structures of cell-cell similarity, without the need for manual specifications of the numbers of DE genes ^11^. (iv) When generating observed counts, we simulate key steps in real experimental protocols, which automatically gives us dropout events, length bias, and distribution of library sizes. We also provide choices to use UMI based protocols or non-UMI full length mRNA protocols, as the properties of data output from these two categories can be very different.

The main input parameters to SymSim are self-explanatory with their own biological or technical meanings, which users can adjust to match an experimental dataset of interest. While the procedure of parameter fitting was developed in order to generate simulated datasets with similar properties, it may also provide additional insight, as the parameters are biologically or technically interpretable. For instance, comparing the parameters fit to the the UMI and non-UMI datasets in this study we note that the capture efficiency inferred to the latter is much higher (Supplementary Material Section 4). The modular nature of SymSim provides possibilities to generalize its application. For example, the generation of true counts with EVFs and transcription kinetics can be replaced by learning a generative model from real data, with methods such as scVI ^4^. This type of extension will facilitate simulation of between-subpopulation diversity that better mimics experimental observations, albeit at the cost of using parameters that are less interpretable biologically. Another extended application of interest is to use different tree structures for different Diff-EVFs when generating multiple populations of cells, such that every tree represents a different aspect of variability between cells. For instance, using this approach, one tree can represent a differentiation process and the other can represent variability due to the physical location of the cell.

As the number and extent of biological applications of single cell genomics continues to grow, so does the extent of analytical questions one can tackle, which go beyond standard ‘bulk era’ analysis steps (e.g., trajectory analysis, mRNA velocity, and more). The need for robust analytical methods therefore increases, and so does the means for proper evaluation of these methods. SymSim provides a starting point to address this challenge of flexible and feature-rich simulation for method evaluation, as it aims to directly mimic the key mechanistic properties of single cell RNA sequencing.

## Methods

### Simulating gene expression with the kinetic model

As shown in Figure 2a, the kinetic model of gene expression considers that a gene can be either *on* or *off* and the probabilities to transit between the two states are *k_on_* and *k_off_*. When the gene is *on* it is transcribed with transcription rate *s*. The transcripts degrade with rate *d*. For a given gene, based on these parameters one can simulate the number of its transcript molecules over time. The theoretical probability distribution can be calculated via the Master Equation ^13, 26^, which is the steady state solution for the kinetic model. Alternatively, the kinetic model can be represented by a Beta-Poisson model, which we use in our implementation to sample expression values for a gene.

### Calculating parameters for the kinetic model in SymSim simulation

For a gene in a cell, the parameters for the kinetic model *k_on_*, *k_off_*, and *s* are calculated from the cell-specific EVF vectors of this cell and the gene effect vectors of the gene (Figure 2a). To allow independent control of the three parameters, we use one EVF vector and one gene effect vector for each parameter. Take *k_on_* as an example: denoting the EVF vector as 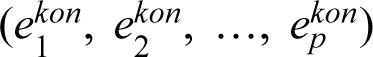, and the gene effect vector for *k_on_* as 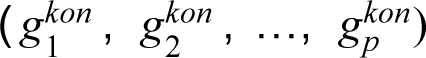, the cell-gene specific value for *k_on_* is the dot product of these two vectors. We then map these *k_on_* values to the distribution of *k_on_* estimated from experimental data, to obtain the matched parameters. We do so by sorting the *k_on_* values (from dot products) for all genes in all cells, sampling the same number of values from the experimental *k_on_* distribution (the number of values would be *m*\**n*, where *m* is the number of genes and *n* is the number of cells), and updating the *k_on_* values to the ones sampled from the experimental distribution with the same rank. The values of *k_off_* and *s* are calculated in the same way.

### Estimating kinetic parameters from real data

We estimated kinetic parameters from experimental data using an MCMC approach. For each gene, its expression *X* depends on *p*, the proportion of time it is *on*, and the mRNA synthesis rate *s*.The parameter *p* itself is a random variable determined by the kinetic parameters *k_on_* and *k_off_*. We model *p* as a Beta distributed variable with shape parameters *k_on_* and *k_off_*. We model X as a Poisson distributed variable with parameter *p*s*. The distribution of X is then identical to the distribution calculated using the Master Equation ^45^. The downsampling effect is modeled as a Binomial sampling with *X* being the number of trials, and *f* being the probability that a transcript is sampled for sequencing.

We fit this model to the experimental data using the Gibbs sampler implemented in RJAGS. At every iteration, we sample each parameter from its marginal posterior conditional on the value of all other parameters. To meet the assumption that all cells share the same kinetic parameters we divide cells by clustering that is performed in the original study and fit the model to counts in a single cluster of cells at a time. We also use imputed read counts, rather than the raw read counts. We use scVI^4^ and ZINB-WaVE ^17^ for the imputation. Since MCMC is dependent on initial conditions, we fit the model independently three times, for each cell cluster and each imputation method. We thinned the MCMC chain to reduce the effect of autocorrelation, and combined all results to obtain the final distribution of kinetic parameters.

### Ranges of kinetic parameters from literature

We look into literature for the ranges of kinetic parameters *k_on_*, *k_off_*, and *s* which are experimentally measured ^18–25^. The range of burst size, or *s*, from these studies ranges from 2-4000. And the *k_on_* and *k_off_* values ranges from 0.0001 to 1 per minute, and the half-life of mRNA varies from 1-10 hours, which correspond to 0.001 to 0.01 per minute. This means that *k_on_*/*d* and *k_off_*/*d* could take values from 10^−2^ to 10^3^. The specific parameter values reported by these studies are in Supplementary Material Section 1.

### Simulation of discrete and continuous populations

The structure of populations can be represented by a tree and the user can input the tree in Newick format in a text file. The differences between populations are realized through Diff-EVFs, which usually account for a small proportion of all EVFs. There are two different modes of simulation the Diff-EVFs, Continuous and Discrete. Both modes can be modeled by Brownian motion along the tree from root to leaves, where one starts with a given value at the root (default is 1), and at each time point *t*, y(t) is calculated as *y*(*t*) = *y*(*t* − 1) + *N* (0, Δ*t*), where *N* () represents a Gaussian function, and Δ*t* is the step size. The values at internal nodes of the tree are shared by all branches connecting this node. In the continuous mode, the step size between two consecutive cells on a given branch is obtained by randomly sampling *n_b_* positions on a branch *b* of length *l_b_*. In the discrete mode, the step size is the corresponding branch length and the number steps is the depth of the tips. For a given tip and a given EVF, the value we sample at the tip is used as the mean of a Gaussian distribution to sample the values for that EVF for all cells in that population with standard deviation σ (Figure 3a).

For the continuous mode we also provide an alternative option to the Brownian motion model: using impulse functions for modeling the path-specific variation. When impulse function is used, for cells sampled from branches that are not on the root-tip path for the specific EVF, they are sampled from a univariate normal with mean equal to the EVF value at their most recent common ancestor with the varying path, and standard deviation *σ*.

### Simulation of technical steps from mRNA capturing to sequencing

We simulate two categories of library preparation protocols, one does not use UMIs (unique molecular identifiers)^34^ and sequences full length mRNAs (using procedures in Smart-seq2 ^33^ as template), and the other uses UMIs and sequences only the 3’ end of the mRNA (using the Chromium chemistry by 10x Genomics as template). In the pre-amplification step, we provide option of using linear amplification to mimic the CEL-seq protocol. As shown in Figure 4a, we take one transcript with 16 molecules as an example. To implement the UMIs, each original molecule has a variable to its count at each step. The technical steps include the following:

1. Capturing step: molecules are captured from the cell with probability α.
2. Pre-amplification step: if using non-CEL-Seq protocols, this step involves N rounds of PCR amplifications. We introduce sequence-specific biases during amplification, which includes transcript length bias and other bias assigned randomly. Parameter *lenslope* can be used to control the amount of length bias, and *MaxAmpBias* is used to tune the total amount of amplification bias. If using CEL-Seq protocol this step is the *in vitro* transcription (IVT) linear amplification.
3. Fragmentation step: the mRNAs are chopped into fragments for sequencing. If sequencing full-length mRNA, all fragments with acceptable length are kept for sequencing. If sequencing only the 3’ end for UMI protocols, only fragments on the 3’ end are kept for sequencing. For each transcript length, we calculate a distribution of number of fragments given expected fragment length, and use this distribution to generate the number of fragments during our simulation of the fragmentation step. The distributions are different for non-UMI and UMI protocols, and the details of calculating the distributions are in Supplementary Material Section 2.
4. Amplification step: fragments go through another k rounds of PCR amplifications for all protocols, including CEL-Seq and non-CEL-Seq protocols.
5. Sequencing step: amplified fragments from the previous step are randomly selected according to a given value of sequencing depth, which is the total number of reads (fragments) to sequence.
6. After the sequencing step (assuming all reads are correctly sequenced and mapped to their original gene), we can get the UMI counts for UMI protocols and read counts for non-UMI protocols.

Note that for simplicity, this pipeline omits several steps, including reverse transcription, and library cleaning up.

### Simulation of amplification biases

During PCR amplification of the full length cDNAs, the PCR amplification rate (namely, probability to be amplified) can vary for different transcripts. As a result, some transcripts are over- and some are under-amplified. This causes the unwanted amplification bias. To simulate this, for each gene, its PCR amplification rate is set to a sum of a basal amplification rate (the input parameter *rate_2PCR*, which equals to the average amplification rate across all genes) plus a bias term *B*. The bias term *B* ranges from -*MaxAmpBias* to *MaxAmpBias*, where *MaxAmpBias* is a user-specified parameter to represent the total amount of amplification bias in our system.

*B* is composed of two categories of biases: biases related to transcript length (referred to as gene length bias, denoted by *B_length_*) and biases caused by other factors (denoted by *B_rand_*). We use a linear function to model the gene length bias: we first bin all gene lengths into *nbins* bins, and get the average length in each bin: *L* = (*l_bin_*_(1)_, *l_bin_*_(2)_,…, *l_bin_*_(*nbins*)_). The length bias term associated with a gene in bin *i* is set to: *B_length_* (*i*) = *lenslope* * *median*(*L*) − *lenslope* * *L*(*i*)

The parameter *lenslope* controls the extent of gene length bias. To ensure that *B_length_* does not exceed *MaxAmpBias*, the parameter *lenslope* should be smaller than 2 × *MaxAmpBias*/(*nbins* − 1). We then set the second term *B_rand_* to a random value in the residual range [-*MaxRandBias, MaxRandBias*] where *MaxRandBias* = *MaxAmpBias* − *max*(*B_length_*). Namely, *B_rand_* = *N* (0, *MaxRandBias*), where N is a Gaussian.

Therefore, for a given gene with length *l*, its PCR amplification rate is:

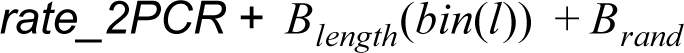

This rate is used in all rounds of PCRs in the pre-amplification step. The biases then get amplified as more PCR cycles are performed, where transcripts with higher amplification rate will likely get more molecules. Assigning a UMI to each molecule before amplification allows us to collapse all molecules with the same UMI after amplification, so different amplification rates will not affect the final molecule counts. For Figure 4b, *lenslope* is set to 0.023, *MaxAmpBias* is set to 0.3, *nbins* is set to 20, and *rate_2PCR* is set to 0.7.

### Fitting simulation parameters to real data

To find the best matching parameters to a real dataset, we simulate a database of datasets with a grid of parameters over a wide range. For each simulated dataset, we calculate the following statistics: mean, percent-non-zero, standard deviation of genes over all cells. Then given a real dataset, we find the simulated dataset which have the most similar distributions of the statistics to the real data, and return the corresponding parameter configurations.

The parameters and their ranges for simulating the two databases are as following:

| parameters | Values for the non-UMI database | Values for the UMI database |
| --- | --- | --- |
| Sigma ($\sigma$) | 0.2, 0.6, 1 | 0.2, 0.6, 1 |
| Alpha_mean ($\alpha$) | 0.05, 0.1, 0.15, 0.2 | 0.01, 0.04, 0.07, 0.1 |
| Alpha_sd ($\beta$) | 0.015, 0.03, 0.045, 0.06 | 0.003, 0.012, 0.021, 0.03 |
| depth_mean (*Depth*) | 1e+05, 5e+05, 1e+06, 2e+06 | 40000, 70000, 1e+05 |
| Depth_sd (*Depth_sd*) | 30000, 150000, 3e+05, 6e+05 | 12000, 21000, 30000 |
| gene_effects_sd | 1, 2 | 1, 2 |
| gene_effect_prob (1-η) | 0.1, 0.3 | 0.1, 0.3 |
| nevf (# EVFs) | 10, 30, 50 | 10, 30, 50 |

### Applying dimensionality reduction and clustering methods

We apply three different dimensionality reduction methods to cluster cells simulated from multiple discrete populations: PCA, scVI and SIMLR. PCA is the naive baseline method that is also the most commonly seen in single-cell RNA-seq analysis. scVI is a more recent method that uses a zero-inflated negative binomial variational auto-encoder model to infer latent space for each single cell. For both the first two methods, cluster identities are then assigned using k-means clustering. The third method, SIMLR, performs dimensionality reduction and cluster identity iteratively to maximize cluster separation.

### Simulation of differentially expressed (DE) genes

Diff-EVFs give rise to differences between populations as well as DE genes between populations. DE genes by design are the ones with non-zero gene effect values corresponding to the Diff-EVFs (Figure 6a), as the gene effect vectors are sparse with a majority of values being 0s. However, the actual expression values of these genes might not differ if the changes in two EVFs cancel out, or if the effect of change one of the kinetic parameter is canceled out by the change in another kinetic parameter. Thus we also use the log fold change of mean gene-expression from the two populations as another criteria. The mean expression can be calculated based on simulated true counts, which is subject to gene-expression intrinsic noise, or based on the kinetic parameters themselves, directly from the theoretical gene-expression distribution. If the kinetic parameters of a gene in a cell is *k_on_*, *k_off_* and *s*, the expected gene-expression of this gene in this cell is *s*\**k_on_*/(*k_on_*+*k_off_*).

### Detection of differentially expressed (DE) genes

DE genes in observed counts are detected respectively with edgeR, DESeq2, Wilcoxon test and Student t-test. For edgeR, we used the quasi-likelihood approach (QLF) with cellular detection rate as covariate. For DESeq2, we use “local” for the fittype parameter, and we evaluate its performance respectively based on the p-values and adjusted p-values which serve as filtering of genes.

### Binomial model to calculate the probability of detecting a population

Assuming all sequenced cells are correctly assigned to its original population, the probability that at least *x* cells are detected from a population only depends on the binomial sampling. Denote the total number of cells by *N* and the proportion of the cells in the given population by *r*, the probability that at least *x* cells are detected for the population is:

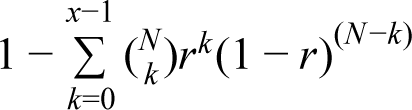

This formula is used to generate the black curves in Figure 7a-d.

### Sources of experimental data

We use two experimental datasets throughout this paper, one is from ^16^ which does not use UMIs and the other is from ^15^ which uses UMIs. The former datasets profiles Th17 cells under various conditions and in our paper we use a subpopulation of 130 TGF-β1+IL-6 cells. We refer to this dataset as the Th17 dataset. The second dataset profiles 3006 cerebral cortex cells. The authors found nine classes in these cells. In this paper, to get distributions of kinetic parameters to map our simulated parameters to the same distribution, we perform parameter estimation respectively on 1) 628 cells sampled from the oligodendrocyte class; 2) 715 cells sampled from the CA1 pyramidal neurons; 3) 296 cells sampled from the S1 pyramidal neurons. To verify that SymSim can simulate data with similar statistics with given experimental dataset, we use all 948 oligodendrocyte cells.

## Acknowledgements

XZ was supported by grant #220558 from the Ragon Institute of MGH, MIT and Harvard. CX and NY were supported by NIH/NHLBI grant U19 AI-090023-09.

## Author contributions

XZ, CX and NY conceived the study and wrote the paper. XZ and CX wrote the software and conducted the analysis.

---

**Figure S1.**
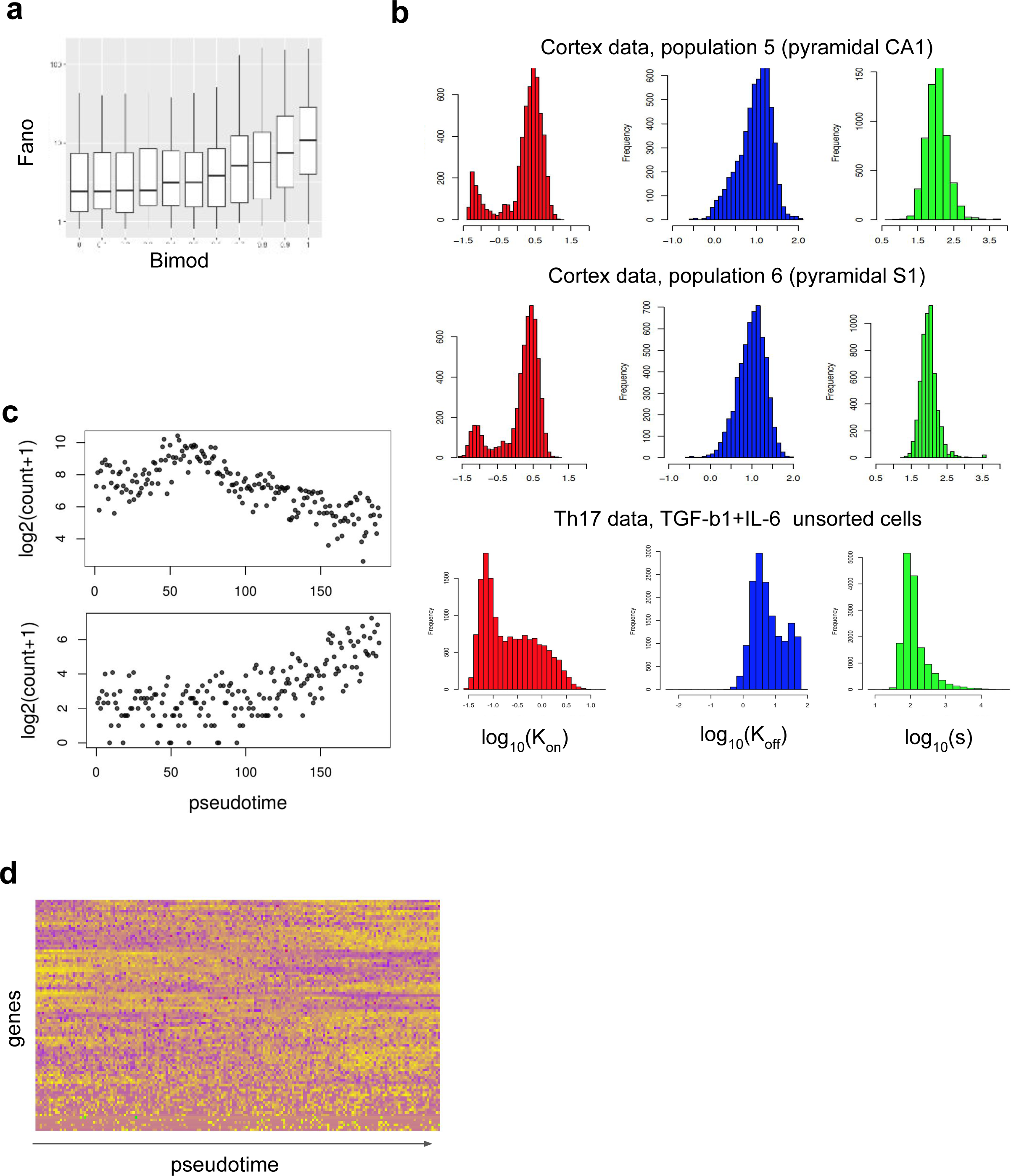
(**a**) The Fano factor of genes across cells with different values for *Bimod*. (**b**) Distribution of kinetic parameters estimated from different experimental data. (**c**) The gene expression of two DE genes along the pseudotime in continuous populations. The structure of populations is represented by the tree in Figure 3. We plot gene-expression of the lineage from the root to population 2. The number of EVFs is 20 for each of the three kinetic parameters, and 12 EVFs of parameter *s* are Diff-EVFs. σ is set to 0.4. (**d**) Heatmap of expression of genes with at least 4 Diff-EVFs along the lineage from the root to population 2.

**Figure S2.**
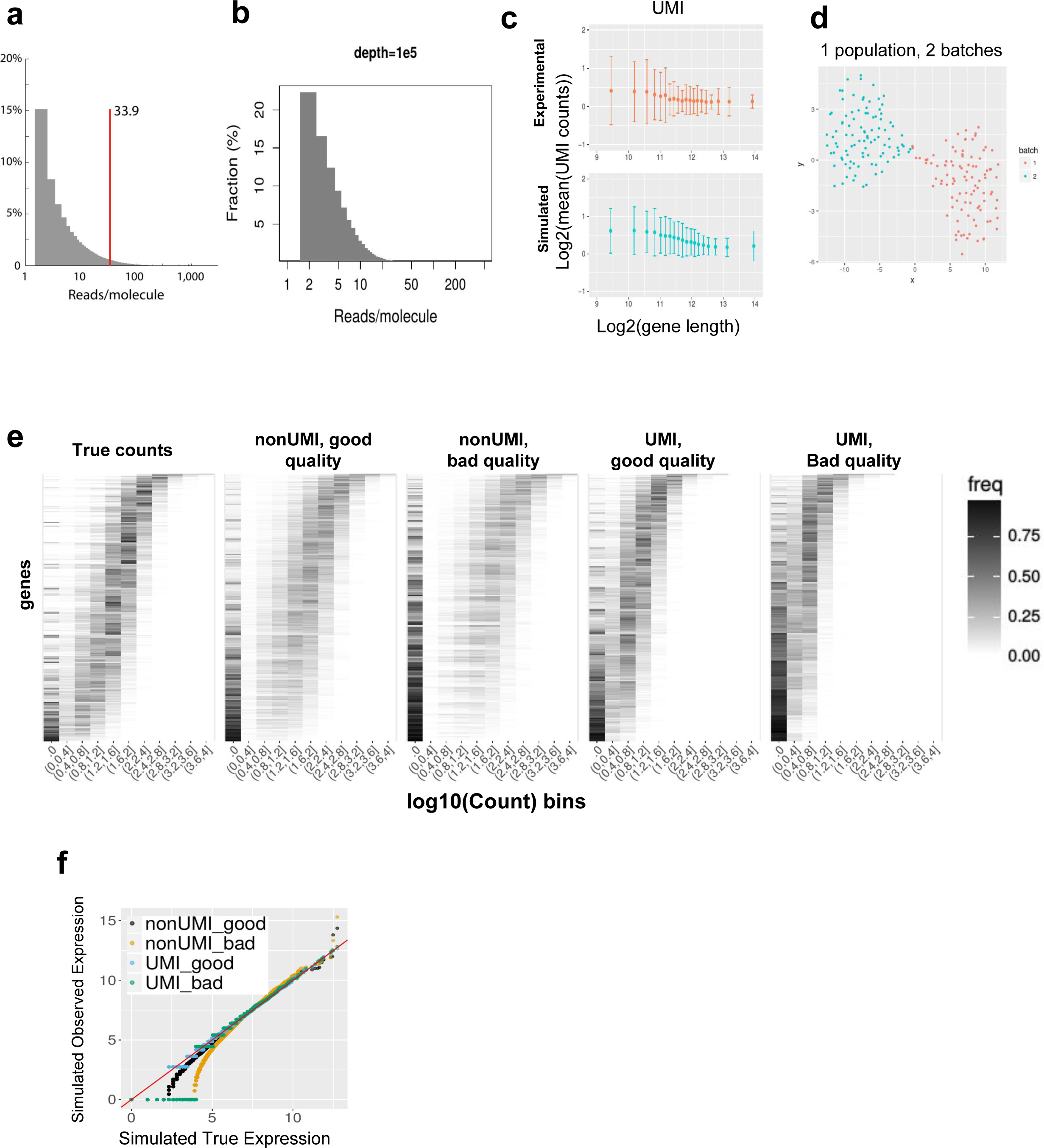
(**a**) The distribution of number of reads per UMI sequenced in the cortex dataset. This plot comes from the supplementary material of the original paper ^15^. (**b**) The distribution of number of reads per UMI sequenced in our simulated data, when using the UMI protocol, with 100k reads per cell. (**c**) The gene length bias in observed counts with UMI protocol, respectively from experimental and simulated data. The experimental data is the cortex dataset from paper^15^. (**d**) TSNE plot of cells simulated for one homogeneous population in two batches. (**e**) The histogram heatmap of gene expression of true simulated counts, and observed simulated counts under “good” and “bad” parameter settings. The parameters are the same as described in Figure 4c. In these heatmaps, each row corresponds to a gene, each column corresponds to a level of expression, and the color intensity is proportional to the number of cells that express the respective gene at the respective expression level. (**f**) Q-Q plots of gene expression of true simulated counts and observed simulated counts under “good” and “bad” parameter settings. The data is the same as that plotted in Figure 4c.

**Figure S3.**
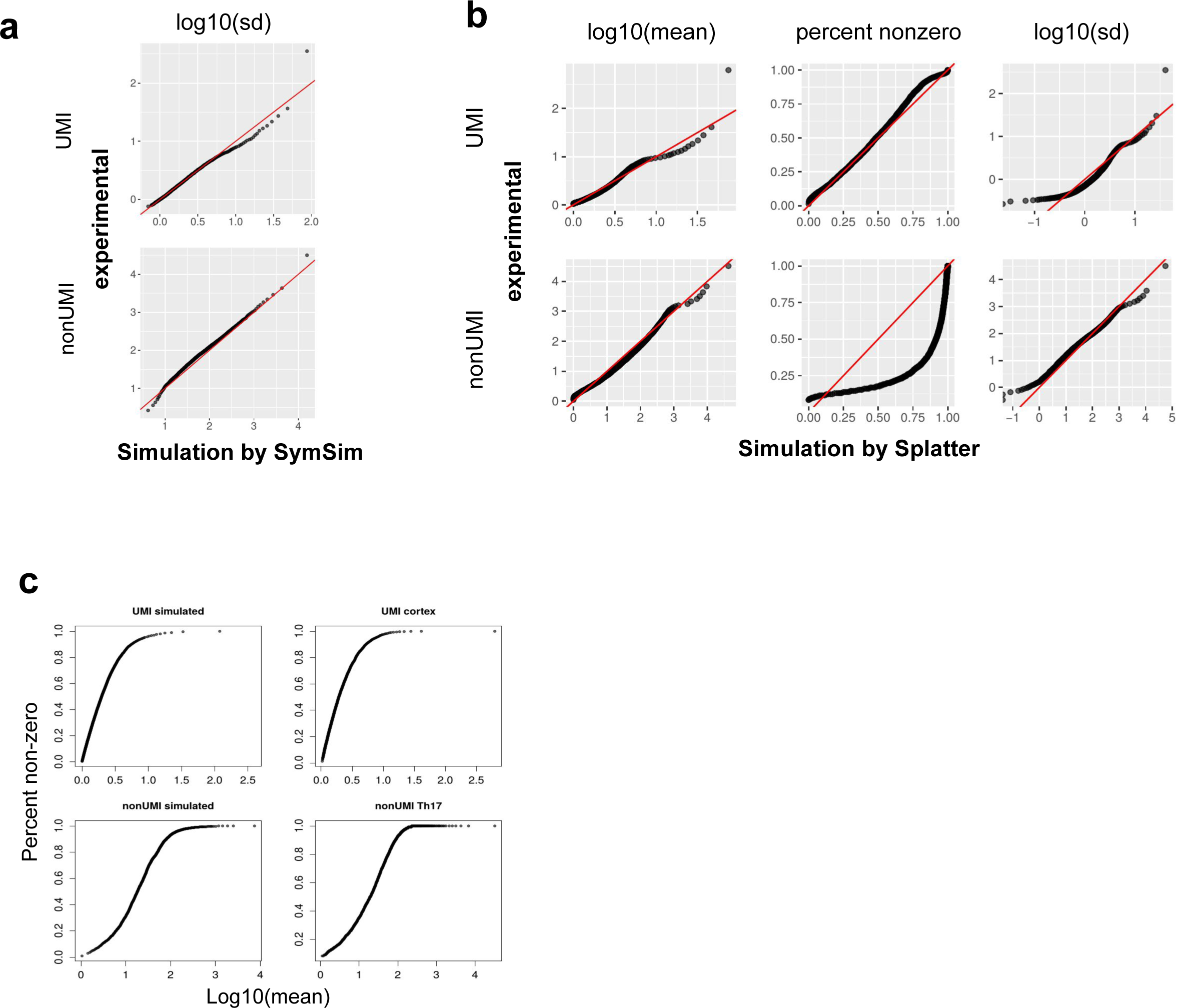
(**a**) Q-Q plots of standard deviation for genes between experimental data and SymSim simulated data for both UMI and non-UMI protocols after removing a proportion of lowly expressed genes. (**b**) Q-Q plots of mean gene expression, percent of non zeros and standard deviation for each gene between UMI and non-UMI experimental data and Splatter simulated data. (**c**) The curve of mean gene expression *vs* percent of non zeros (detection rate) in simulated UMI data, experimental UMI data (the cortex dataset), simulated non-UMI data, experimental non-UMI data (the Th17 dataset).

**Figure S4.**
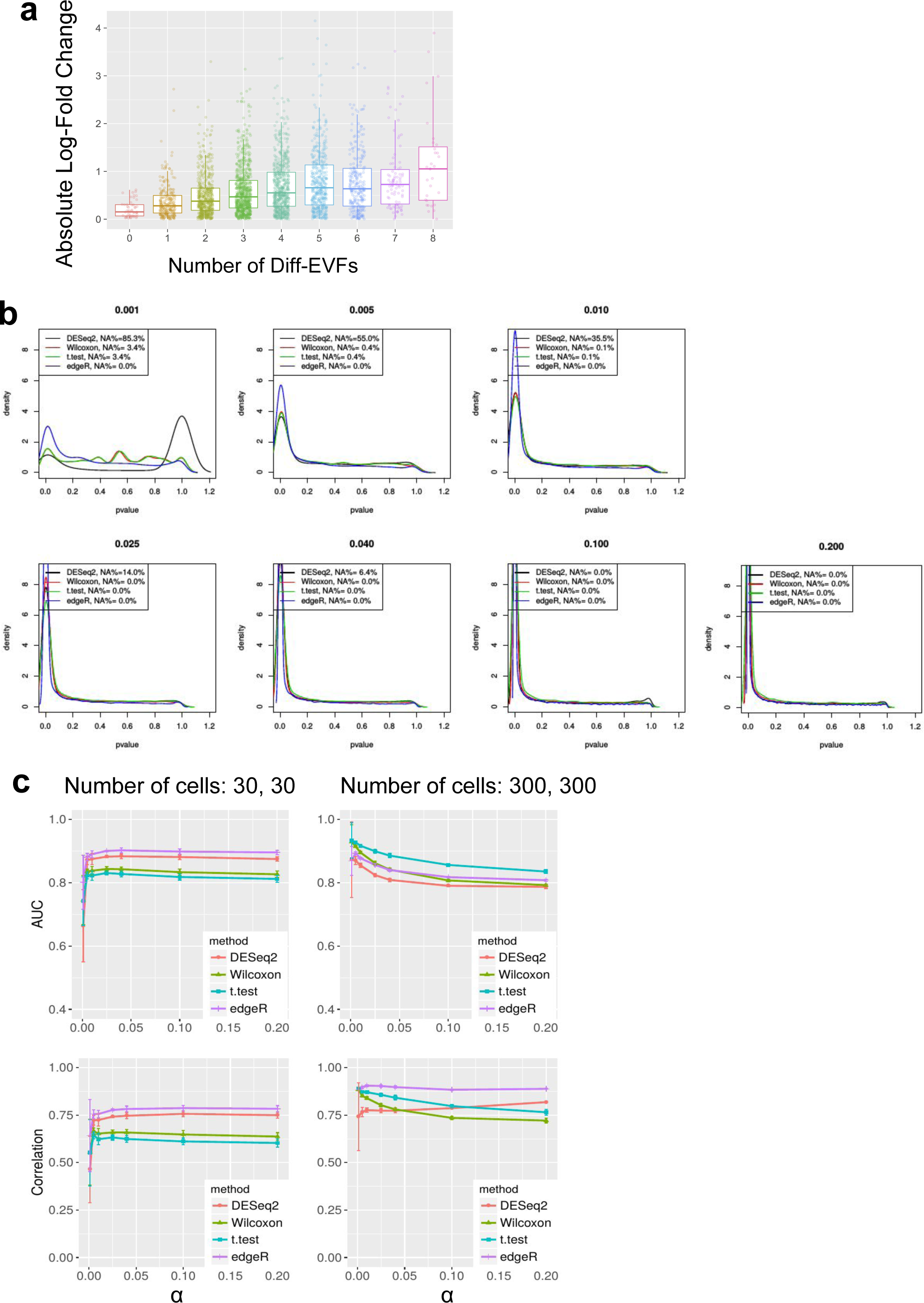
(**a**) The log fold change of genes with different number of Diff-EVFs. (**b**) The distribution of p-values of four different DE methods, with different values of capture efficiency α (noted as the title of each plot). (**c**) The AUROC (top row) and negative of correlation (bottom row) measures of the four different methods for DE gene detection. In these plots the adjusted p-values from DESeq2 are used instead of the p-values, and the genes with NAs in the adjusted p-values are removed for all methods to calculate the accuracy measures. The numbers of cells in both populations for the plots on the left column are both 30, and on the right column the numbers of cells are both 300.

**Figure S5.**
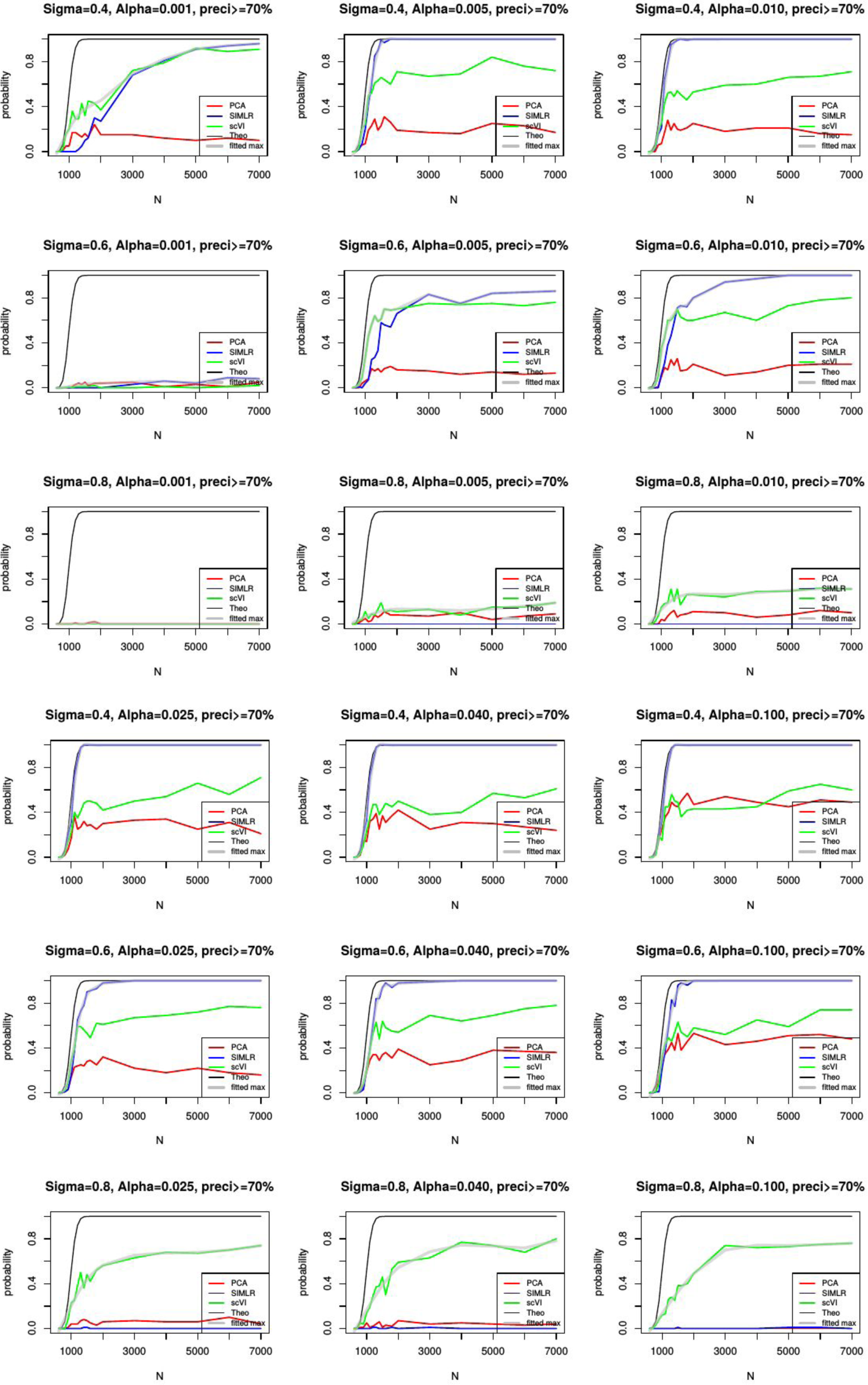
The probability of detecting the rare population (population 2 in the tree shown in Figure 3) under a wide range of configurations of *σ* (Sigma) and *α* (Alpha). The criteria of detecting the population is that at least 50 cells are detected (true positive >=50) with precision at least 70%.

