## Supplementary Material for "SymSim: simulating multi-faceted variability in single cell RNA sequencing"

Supplementary Materials

#### 1. Kinetic parameters from experiments

| Paper | Genes | Method | Parameters |
|---|---|---|---|
| Padovan-Merhar, O. and Raj, A. 2015. Single Mammalian Cells Compensate for Differences in Cellular Volume and DNA Copy Number through Independent Global Transcriptional Mechanisms. Molecular Cell. 58, 2 (2015), 339–352. | UBC, MYC, EEF2, TUSC3 | smFISH<br><br>Transcriptional Blockage | Burst size 500-4000 (mean transcription site intensity)<br>On frequency 20-75% ( number of transcription site per DNA copy)<br>Degradation rate per volume 0.25-0.5 |
| Bahar Halpern, K. et al. 2015 Bursty gene expression in the intact mammalian liver. Molecular cell. 58, 1 (Apr. 2015), 147–56. | Achy, Actb, Ass1, Fasn, G6pc, Pck1, Srebf1,Insr | smFISH | burst size 1-700 $hr^{-1}$<br>kon 0.01-1.32 $hr^{-1}$<br>koff 0.021-3.36 $hr^{-1}$ |
| Skinner, SO et al. 2016. Single-cell analysis of transcription kinetics across the cell cycle. Elife. (2016). | Oc4,Nanog | smFISH | $k_{on} \approx 9\times10^{-3}$ $min^{-1}$ for Oct4, $2\times10^{-3}$ $min^{-1}$ for Nanog<br>$k_{off} \approx 2\times10^{-2}$ $min^{-1}$ for Oct4, $7\times10^{-3}$ $min^{-1}$ for Nanog<br>34% on for Oct4, 22% on for Nanog<br>$S\approx$0-6 mRNA per site for Oct4, 0-4 mRNA per site for Nanog |
| Dey, Siddharth S., et al. "Orthogonal control of expression mean and variance by epigenetic features at | HIV LTR | smFISH | $S/k_{off}$ (Burst size): 2-24<br>$k_{on}/d$ (Burst frequency): 0.3-4.5 |

| | | | |
|---|---|---|---|
| different genomic loci." Molecular systems biology 11.5 (2015): 806. | | | |
| Singh, Abhyudai, et al. "Dynamics of protein noise can distinguish between alternate sources of gene-expression variability." Molecular systems biology8.1 (2012): 607. | HIV LTR | Transcriptional blockage, smFISH | $S/k_{off}$ (Burst size): 2-12<br>Degradation rate is reported in fluorescent density, thus is not comparable to other results) |
| Xu, H et al. 2015. Combining protein and mRNA quantification to decipher transcriptional regulation. Nature methods. (2015). | *hunchback* | smFISH | $k_{on}$ and $k_{off}$ from 0 to 10 $min^{-1}$<br>$k_{INI}$ from 0 to 100 $min^{-1}$<br>$S$ from 0-100 $min^{-1}$ |
| Suter, D.M. et al. 2011. Mammalian genes are transcribed with widely different bursting kinetics. Science (New York, N.Y.). 332, 6028 (Apr. 2011), 472–4. | Bmal1a, Glutaminase, Prl2C2 | smFISH | kon 0.02-0.06 min-1<br>koff 0.1-0.6 min-1<br>s 3-18 min-1 |
| Bartman et al. 2016. Enhancer regulation of transcriptional bursting parameters revealed by forced chromatin looping. (2016). | beta-globin | smFISH | Proportion on: 0.25-0.75 |
| Rabani, Michal, et al. "A Massively Parallel Reporter Assay of 3′ UTR Sequences Identifies In Vivo Rules for mRNA Degradation." | 3' UTR sequences during early zebrafish embryogenesis | MPRA | half-life of mRNA is 1-10h (d = 0.01155245 min-1 ~ 0.001 min-1) |
| Hendy, Oliver, et al. "Differential context-specific impact of individual core promoter elements on transcriptional dynamics." Molecular biology of the cell 28.23 (2017): 3360-3370. | MHC class 1 genes | smFISH | $S/k_{off}$ (Burst size): 5.5-13.9<br>$k_{on}/d$ (Burst frequency): 4.4-13.9<br>Proportion on: 0.15-0.71 |

#### 2. Setting branch lengths and σ to get clusters of different clusterability

In the function of generating multiple discrete populations, users can control the extent of between-population variation by setting the branch lengths of the input tree, and control the within-population variation by parameter $\sigma$. Notably, both $\sigma$ and the square root of branch lengths in the tree are in units of EVF values. For any given Diff-EVF and any two given populations, the ratio of square root tree distance to $\sigma$ determines the overlap between the distributions of the two Diff-EVFs. Thus this ratio determines the separability between the two populations. Take the Diff-EVF1 of populations 2 and 3 in Figure 3 as an example: we can show that

$$E(|y_1(1) - y_2(1)|) = \sqrt{d_{23}} \cdot \sqrt{2/\pi} \qquad \text{(Equation 1)}$$

where $d_{23}$ is the distance in the tree between Populations 2 and 3. As the EVF values of Diff-EVF1 for cells in Populations 2 and 3 are sampled respectively from distributions $N(y_1(1),\sigma^2)$ and $N(y_1(2),\sigma^2)$ (Figure 3), the ratio $H = \frac{E(|y_1(1) - y_2(1)|)}{\sigma} = \frac{\sqrt{d_{23}} \cdot \sqrt{2/\pi}}{\sigma}$ correlates with the separability between cells from Population 2 and cells from Population 3. Detailed derivation and proof of Equation 1 are in Supplementary File 1.

#### 3. Distributions of number of fragments

When simulating the fragmentation step, we need the number of fragments obtained from a transcript. This number is dependent on the transcript length (denoted by *L*), the read length (*r*), maximum fragment length (*f*) and expected gap size (*g*) of the reads assuming we use paired-end sequencing. The fragmentation efficiency which is the probability with which a cut happens to a position on the transcript is: $e=1/(2*r+g)$.

For nonUMI protocols where full length mRNA is sequenced, for each transcript length, we simulate the fragmentation process many times with the probability *e* and remove resulting pieces which have length smaller than *r* or greater than *f*, and we obtain a distribution of number of valid fragments for a given transcript length. In SymSim, we just sample from this distribution.

For UMI protocols, we only need the valid fragments at the 3' end. In this case, we can derive theoretical distributions of the probability that a mRNA copy gives rise to a fragment. The expressions are as follows:

$(1-e)^r(1-(1-e)^{(f-r-1)})$ , if $L \geq f$

$(1-e)^r$ , if $r < L < f$

$0$ , if $L \leq r$

So during SymSim we sample with these probabilities to get the number of 3' end fragments (which will be either 0 or 1).

In our paper, we set *r*=100, *f*=1000, *g*=200.

### 4. Top parameters which give rise to simulated datasets similar to real data sets

With SymSim we generate a database where datasets are simulated with a large grid of parameters. We also calculate a "summary" for the dataset corresponding to each parameter configuration. This summary includes mean expression, percentage of expressing cells, and Fano factor for each gene. Given an experimental dataset, one can use these statistics to find the best matching simulations in our database. The parameters which yield the best matching simulations can give us insights on the properties of the experimental dataset. In Table 1 we show the top 8 parameter configurations which match best to population 3 in the Cortex dataset (UMI), and in Table we show the top 8 configurations for the Th17 dataset (nonUMI). For both datasets, the top 8 parameters give very consistent values for Sigma, which denotes the heterogeneity of the cells; alpha_mean, which is the mean capture efficiency; and depth_mean, which is the mean sequencing depth. We see that the Th17 dataset has higher capture efficiency and sequencing depth than the cortex dataset, and more homogeneous (lower $\sigma$).

**Table 1 Top parameters which match the Cotex UMI dataset**

| Gene_effects_sd (standard deviation to sample gene effect values) | gene_effect_prob (1-η) | nevf | Sigma ($\sigma$) | Alpha_mean ($\alpha$) | Alpha_sd ($\beta$) | Depth_mean (*Depth*) | Depth_sd (*Depth_sd*) |
|---|---|---|---|---|---|---|---|
| 1 | 0.3 | 10 | 0.6 | 0.04 | 0.012 | 1e5 | 30000 |
| 2 | 0.1 | 30 | 0.6 | 0.04 | 0.012 | 1e5 | 30000 |
| 2 | 0.1 | 50 | 0.6 | 0.04 | 0.012 | 1e5 | 30000 |
| 1 | 0.1 | 30 | 0.6 | 0.04 | 0.012 | 1e5 | 30000 |

| | | | | | | | |
|---|---|---|---|---|---|---|---|
| 1 | 0.1 | 50 | 0.6 | 0.04 | 0.012 | 1e5 | 30000 |
| 2 | 0.3 | 50 | 0.6 | 0.04 | 0.012 | 1e5 | 30000 |
| 2 | 0.3 | 30 | 0.6 | 0.04 | 0.012 | 1e5 | 30000 |
| 2 | 0.3 | 10 | 0.6 | 0.04 | 0.012 | 1e5 | 30000 |

**Table 2 Top parameters which match the nonUMI Th17 dataset**

| Gene_effects_sd (standard deviation to sample gene effect values) | gene_effect_prob (1-η) | nevf | Sigma ($\sigma$) | Alpha_mean ($\alpha$) | Alpha_sd ($\beta$) | Depth_mean (*Depth*) | Depth_sd (*Depth_sd*) |
|---|---|---|---|---|---|---|---|
| 2 | 0.3 | 10 | 0.2 | 0.1 | 0.03 | 2e6 | 6e5 |
| 2 | 0.1 | 50 | 0.2 | 0.1 | 0.03 | 2e6 | 6e5 |
| 1 | 0.1 | 50 | 0.2 | 0.1 | 0.03 | 2e6 | 6e5 |
| 1 | 0.3 | 30 | 0.2 | 0.1 | 0.03 | 2e6 | 6e5 |
| 2 | 0.1 | 30 | 0.2 | 0.1 | 0.03 | 2e6 | 6e5 |
| 1 | 0.1 | 10 | 0.2 | 0.1 | 0.03 | 2e6 | 6e5 |
| 2 | 0.3 | 50 | 0.2 | 0.1 | 0.03 | 2e6 | 6e5 |
| 1 | 0.1 | 30 | 0.2 | 0.1 | 0.03 | 2e6 | 6e5 |

Other parameters we keep fixed for all UMI and non-UMI datasets are:
evf_center=1, geffect_mean=0, bimod=0, lenslope=0.01, nbins=20, MaxAmpBias=0.2, rate_2PCR=0.8, nPCR=18.

### 5. Parameter settings and measurements for benchmarking and experimental design

5.1 Parameters used for length bias in Figure 4b

Key parameters which give rise to the length bias patterns shown in Figure 4b are: $\alpha$=0.05, *lenslope*=0.023, nbins=20, *MaxAmpBias*=0.3, *Depth*=1.3e6.

5.2 Parameters used for regression in Figure 5a

| Parameter | Values | # simulations per value |
| --- | --- | --- |
| N (total number of cells) | 1000 2000 4000 6000 8000 | 1440 |
| Prop (proportion of rare population) | 0.01 0.03 0.05 0.10 0.20 | 1440 |
| σ (within population variability) | 0.4 0.6 0.8 1.0 | 1800 |
| α (capture efficiency) | 0.001 0.005 0.010 0.025 0.050 0.100 | 1200 |
| Depth (sequencing depth) | 5e+03 1e+04 5e+04 1e+05 | 1800 |

We have kept the standard deviation (SD) of α and Depth small for all simulations so that the SD do not dominate the mean in cases of low α and Depth values. The SD for α is 5e-04, and SD for Depth is 1000.

5.3 Parameters used for benchmarking of DE methods in Figure 6d and for “How Many Cells” analysis in Figure 7

| Parameter | Values |
| --- | --- |
| Prop (proportion of rare population) | 0.2 |
| n_de_evf (# Diff-EVFs for *s* only) | 18 |
| σ (within population variability) | 0.4 0.6 0.8 1.0 |
| α (capture efficiency) | 0.001 0.005 0.010 0.025 0.05 0.1 0.2 |
| Depth (sequencing depth) | 1e+05 |
