## Supplementary File 1 for "SymSim: simulating multi-faceted variability in single cell RNA sequencing"

### Relationship Between Tree Branch Length and Sigma

Chenling Xu

March 22, 2018

1. Diagram of the tree
   $R$: EVF value at the root of the tree, for simplicity R=0
   $W, X, Y$: EVF value at the tips of the tree
   $Z$: EVF value at the most recent common ancestor of two populations.
   $a, b, c, d$: Branch length
   Let EVF values perform Brownian Motion with constant rate 1 for time equal to the branch length along the tree. The random variables at the tip of the tree represent the EVF population mean.

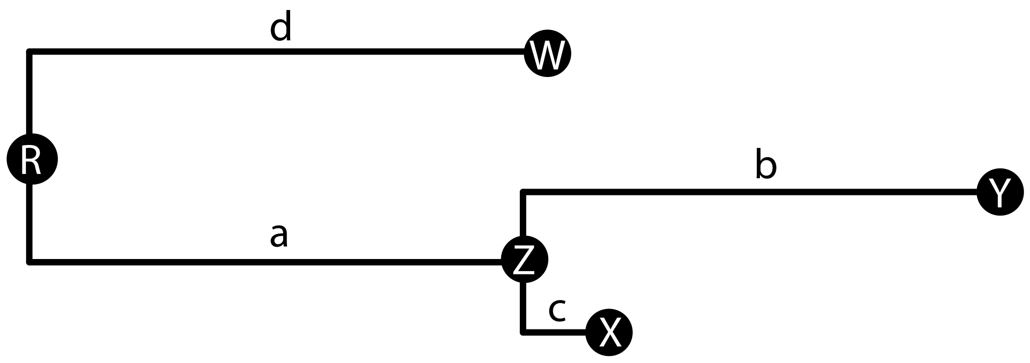


2. Relationship between Branch Length and EVF-Mean Variance-Covariance Matrix

(a) EVF values at the tips of the tree are normally distributed with mean $\mu$ and variance equal to the sum of branch lengths between the tips and the root, with $\mu$ being the value at the root.
Without loss of generosity, let $\mu = 0$

$$W \sim N(0, d)$$

We can reformulate Brownian Motion in terms of a 1 Dimensional Random Walk. In each unit time, EVF values take $k$ steps of size $\frac{1}{\sqrt{k}}$ in random directions. Let the direction of the step be $\delta$ and it can take either value 1 or -1.

$$W = \frac{1}{\sqrt{k}} \sum_{i=1}^{dk} \delta_i$$

This can be re-written as

$$W = \sqrt{d}(\frac{1}{\sqrt{dk}} \sum_{i=1}^{dk} \delta_i)$$

Because $\delta_i$ are i.i.d with mean 0 and variance 1, by central limit theorem their normalized sum $\frac{1}{\sqrt{dk}} \sum_{i=1}^{dk} \delta_i$ has standard normal distribution. Thus

$$W \sim N(0, d)$$

When $\mu \neq 0$, $W = \mu + \frac{1}{\sqrt{k}} \sum_{i=1}^{dk} \delta_i$, so $W \sim N(\mu, d)$

(b) Covariance of the EVF values at the tips of the tree are equal to the branch length between the root and their most recent common ancestor.

$$Cov(X, Y) = E[(X - E[X])(Y - E[Y])]$$

$$= E[XY - XE[Y] - YE[X] + E[X]E[Y]]$$

Given the last result, we know that $E[X] = E[Y] = 0$

$$Cov(X, Y) = E[XY]$$

$$E[XY] = E[(Z + (Y - Z))(Z + (X - Z))]$$
$$= E[Z^2 + Z(X - Z) + Z(Y - Z) + (X - Z)(Y - Z)]$$
$$= E[Z^2] + E[Z(X - Z)] + E[Z(Y - Z)] + E[(X - Z)(Y - Z)]$$

(c) Claim: Expectation of the product of two independent normal variable with expectation 0 is also zero.
Proof: Let X and Y be two independent normal variable with mean 0 and standard deviation $\sigma_x$, $\sigma_y$. The expectation of their product can be written as

$$E[XY] = \int_{-\infty}^{\infty} \int_{-\infty}^{\infty} XYf(X, Y)dXdY$$

Because of independence,

$$E[XY] = \int_{-\infty}^{\infty}\int_{-\infty}^{\infty} XYf(X)f(Y)dXdY$$

By moving all terms independent of X outside of the inner integral,

$$= \int_{-\infty}^{\infty} Yf(Y)\int_{-\infty}^{\infty} Xf(X)dXdY$$

We know that the expection of X is equal to 0, so

$$\int_{-\infty}^{\infty} Xf(X)dX = 0$$

$$\int_{-\infty}^{\infty} Yf(Y) * 0 \; dY = 0$$

Thus $E[XY] = 0$. Due to the memoriless property of random walk, $Z, (X-Z)$, $Z, (Y-Z)$ and $(Y-Z), (X-Z)$ are all independent. By the property shown in part a

$$E[Z] = E[X-Z] = E[X-Z] = 0$$

thus

$$E[XY] = E[Z^2] = E[Z^2] - E[X]^2 = Var[Z] = a$$

3. Average Distance between EVF-Mean in Two Populations Separated
The average distance, or absolute differences between two populations can be expressed as the $E[|X-Y|]$. It can then be written as

$$E[|(X-Z)+(Z-Y)|]$$

From proof 1a, we know that $X|Z \sim N(Z,c)$,$Y|Z \sim N(Z,b)$,so $(X-Z)|Z \sim N(0,c)$,$(Z-Y)|Z \sim N(0,b)$. Because $X-Z|Z$ and $Z-Y|Z$ are independent, the distribution of their difference given Z is $N(0,b+c)$
Claim: The sum between two independent normal variables $X$ and $Y$ is normally distributed with variance $\sigma_X^2+\sigma_Y^2$. We can show this using characteristic functions.

$$\varphi_X(t) = \mathrm{E}\left(e^{itX}\right), \qquad \varphi_Y(t) = \mathrm{E}\left(e^{itY}\right)$$

By indepence we have

$$\varphi_{X+Y}(t) = \mathrm{E}\left(e^{it(X+Y)}\right)$$

$$\varphi_{X+Y}(t) = \varphi_X(t)\varphi_Y(t) = \exp\left(it\mu_X + \frac{\sigma_X^2 i^2 t^2}{2}\right)\exp\left(it\mu_Y + \frac{\sigma_Y^2 i^2 t^2}{2}\right)$$

$$= \exp\left(it(\mu_X+\mu_Y) + \frac{(\sigma_X^2+\sigma_Y^2)i^2t^2}{2}\right).$$

Which is the characteristic function of a normal distribution with mean $\mu_X + \mu_Y$ and variance $\sigma_X^2 + \sigma_Y^2$. Since we care about the absolute value of their differences, we then need to calculate the absolute value of a normally distributed variable. For simplicity we write the integration for a normally distributed variable $R \sim N(0, \sigma^2)$

$$E[|R|] = \int_{-\infty}^{\infty} |R| f(R) dR$$

$$= \int_{-\infty}^{0} -R f(R) dR + \int_{0}^{\infty} R f(R) dR$$

The density function of a normally distributed variable X is

$$f(X) = \frac{1}{\sqrt{2\pi\sigma^2}} e^{-\frac{(X-\mu)^2}{2\sigma^2}}$$

$$E[|R|] = \int_{-\infty}^{0} -R \frac{1}{\sqrt{2\pi\sigma^2}} e^{-\frac{R^2}{2\sigma^2}} dR + \int_{0}^{\infty} R \frac{1}{\sqrt{2\pi\sigma^2}} e^{-\frac{R^2}{2\sigma^2}} dR$$

Define new variable $u = \frac{-R^2}{2\sigma^2}$. We then have

$$R = \sqrt{-2u\sigma^2}, \frac{dR}{du} = \frac{\sigma^2}{\sqrt{-2u\sigma^2}}$$

Thus

$$\int_{-\infty}^{0} -R \frac{1}{\sqrt{2\pi\sigma^2}} e^{-\frac{R^2}{2\sigma^2}} dR = \frac{1}{\sqrt{2\pi\sigma^2}} \int_{-\infty}^{0} -Re^{-\frac{R^2}{2\sigma^2}} du \frac{dR}{du}$$

$$= \frac{1}{\sqrt{2\pi\sigma^2}} \int_{-\infty}^{0} -\sqrt{-2u\sigma^2} e^{u} \frac{\sigma^2}{\sqrt{-2u\sigma^2}} du$$

$$= \sqrt{\frac{\sigma^2}{-2\pi}} \int_{-\infty}^{0} e^{u} = \sqrt{\frac{\sigma^2}{-2\pi}}$$

Similarly,

$$\int_{0}^{\infty} R \frac{1}{\sqrt{2\pi\sigma^2}} e^{-\frac{R^2}{2\sigma^2}} dR = -\sqrt{\frac{\sigma^2}{-2\pi}} \int_{0}^{\infty} e^{u} = \sqrt{\frac{\sigma^2}{-2\pi}}$$

Thus, $E[|R|] = \sqrt{\frac{2\sigma^2}{\pi}}$

Up to here, we have proven that

$$E[|X - Y||Z] = \sqrt{\frac{2(b+c)}{\pi}}$$

Becuase this expression do not depend on Z,

$$E[X-Y] = \int_{-\infty}^{\infty} f(Z)E[|X-Y||Z]dZ$$

$$= E[|X-Y||Z]\int_{-\infty}^{\infty} f(Z)dZ$$

$$= E[|X-Y||Z] = \sqrt{\frac{2(b+c)}{\pi}}$$

4. The probability of overlap of EVF distribution between two populations
   We use this result to calibrate the value of $\sigma_{within}$ (within population variation) used in our simulation. The average distance of the per-cell EVF to the population mean EVF is equal to $\sqrt{\frac{2\sigma_{within}}{\pi}}$. In our simulation, $\sigma_{within}$ is the same for each population. We can derive the amount of overlap as a function of the distance between the EVE mean and the amount of within population variation.
   For example, if $\sigma_{within} = \frac{a}{2}$, $a$ being the distance between the population mean. We can solve for the point of intersetion of the two density functions

   $$\frac{1}{\sqrt{2\pi\sigma^2}} e^{\frac{-x^2}{2\sigma^2}} = \frac{1}{\sqrt{2\pi\sigma^2}} e^{\frac{-(x-a)^2}{2\sigma^2}}$$

   When we solve for $x$, we see that the two density function intersects at $\frac{a}{2} = \sigma$ from the population means. For normal distributions, the probability of having values greater than $\mu + \sigma$ or smaller than $\mu - \sigma$ is 0.159. Thus the total amount of overlap between the EVF values of the two populations is 2*0.159=0.318. We can generalize this result to other values of $\sigma$ because as long as the value of $\sigma$ is equal for each population, the point of intersection is always the mean of the two population mean. The probability of overlap is then

   $$p(X > \mu + \frac{a}{2}) + p(X \le \mu - \frac{a}{2})$$
